# Phase separation of Polycomb-like (PCL) proteins drive PRC2 complex condensates to regulate gene expression

**DOI:** 10.1101/2025.03.26.645482

**Authors:** Shuangzhou Peng, Bin Zhang, Le Jin, Huixin Xu, Bastian Stielow, Merle Geller, Chuanying Wang, Mengyu Zhang, Xiang Luo, Xijiao Wang, Meizhi Jiang, Xiang Gao, Xiaokun Zhang, Qixu Cai, Fei Sun, Robert Liefke, Rui Gao, Siming Chen

## Abstract

Polycomb group (PcG) proteins function as two major multicomponent protein complexes: Polycomb repressive complex 1 (PRC1) and 2 (PRC2) to repress key developmental genes and maintain epigenetic memory of cell identity during development. Phase-separation has long been implicated in PRC1 mediated gene silencing, but it is still unclear whether PRC2 also utilize a similar mechanism to regulate gene expression. Here we report that Polycomb-like (PCL) proteins, PRC2 accessory factors, can phase-separate *in vitro* and generate dynamic puncta *in vivo*. Biochemical and cellular analyses reveal that PCL proteins (hereafter referred to as PCLs) have intramolecular interaction between N- and C-terminal domains, which make PCLs into more compact conformations. The intramolecular interaction not only controls the size of the PCLs phase separation droplets, but also affects the chromatin association of PRC2. Finally, we show that CpG islands, key DNA regulatory elements in mammalian promoters, disrupt the N-C intramolecular interaction of PCLs to expose their middle intrinsically disordered regions (IDRs), which in turn trigger PCLs driven PRC2 condensates formation. Together, this study provides a new perspective on the regulation of PRC2 by PCLs, implicating PCLs ‘read’ CpG islands to fine-tune their oligomerization to overcome the threshold for PRC2 recruitment to corresponding genomic loci via multivalent interaction with CpG islands chromatin.

## Introduction

Polycomb group (PcG) proteins were first identified in *Drosophila melanogaster* to control body plan segmentation by regulating Hox genes expression^1, 2, 3^. PcG are families of chromatin-modifying protein complexes that transcriptionally silence genes expression by modifying histone tail and partially regulating chromatin structure^4^. PcG proteins function as two main Polycomb repressive complexes (PRCs): Polycomb repressive complexes 1 (PRC1) and 2 (PRC2). While PRC1 mono-ubiquitylates histone H2A on lysine 119 (H2AK119ub1), PRC2 catalyzes mono-, di-, and tri-methylation of lysine 27 on histone H3 (H3K27me1, H3K27me2, and H3K27me3)^5^. Extensive studies demonstrate PRC1 catalyzes H2AK119ub1 and compacts chromatin to silence genes expression likely via liquid-liquid phase separation (LLPS)^6, 7, 8, 9^. PRC1 and PRC2 generally co-occupy silenced gene loci, associate within Polycomb bodies, and coregulate their target genes^10, 11^. Despite this fact, whether the phase separation is also involved in PRC2 mediated gene silencing is still unclear.

Polycomb repressive complex 2 (PRC2) mediated gene silencing is involved in several biological processes including: cell fate specification, X chromosome inactivation, and tumorigenesis^12^. The PRC2 core complex is composed of EZH1/2, EED, SUZ12, and RBBP4/7, which is sufficient to induce H3K27me3 *in vitro*. Proteomics studies have shown that PRC2 core complex can co-purify with a number of accessory factors such as AEBP2, JARID2, PCL1-3, EPOP (also referred to as C17ORF96), and PALI1/2 (also known as C10ORF12), to form different classes of PRC2 holo complexes^13, 14, 15^. These accessory factors either modulate PRC2 catalytic activity or help to recruit PRC2 to appropriate genomic loci.

Recent progress in structural biology has greatly advanced our understanding of function and regulation of PRC2. We and others previous structures of the PRC2 subcomplex and core complex bound to JARID2 and AEBP2 reveal that SUZ12 is architectural element involved in establishing the different PRC2 holo complexes^16, 17^: the C-terminal VEFS domain of SUZ12 together with EZH2 and EED integrate into the minimal catalytic module^18, 19, 20^; the N-terminal region of SUZ12, which scatters on the surface of the RBBP4, provides binding surfaces for JARID2 and AEBP2, which also can be recognized and bound by EPOP and PCLs, respectively^16^. It is worth noting that the structures of PCLs or AEBP2-containing PRC2 adopt a dimeric or monomeric structural architecture respectively^17, 21^, suggesting the oligomeric states of PRC2 complexes can be regulated by different PRC2 accessory factors. A recent cryo-EM structures of PRC2:EZH1 bound to nucleosome demonstrated that PRC2:EZH1 dimers are more effective than monomers at compacting chromatin^22^, allowing us to start to think the possible correlation between phase separation-related chromatin compaction and PRC2 mediated gene silencing. Although PRC2 form microscopically visible condensates that has already been observed in living cells^23^, the mechanisms about regulation of higher oligomeric states of PRC2 are currently unknown.

In humans three PCL family proteins have been identified, PCL1, PCL2, and PCL3 (also known as PHF1, MTF2, and PHF19, respectively)^24, 25^. The PCL1-3 proteins all contain a single Tudor domain followed by two adjacent plant homeodomain (PHD) fingers and an extended homologous (EH) region at the N terminus^26^, and a “reversed chromodomain’’ (RC) located at the C terminus^21^. The Tudor domain mediates the intrusion of PRC2 into active chromatin regions to silence expressed genes by binding to H3K36me3 chromatin modification^27, 28^. The PHD fingers of PHF1 has been shown to be ‘readers’ of H4R3me2s and cooperates with PRMT5 to repress genes expression^29^. The EH domain recognizes DNA and links PRC2 to the CpG islands chromatin^26^. Our previous structure of PCL3/PHF19 containing PRC2 subcomplex suggested that the C-terminal RC region of PCL3/PHF19 stabilizes the dimerization of PRC2 and promotes PRC2 targeting to CpG islands chromatin *in vivo*^21^. In contrast to the extensive effort on the structural and functional studies of the Tudor, PHD fingers, EH domain, and RC region of PCLs, the role of the middle uncharacterized disordered regions (IDRs) of PCLs in regulating PRC2 is largely unknown.

In this study, we show that the PCLs contain middle intrinsically disordered regions (IDRs) that drive biomolecular condensates formation through LLPS. Biochemical and cellular data show that the IDRs driven phase separation are controlled by the intramolecular interaction of PCLs, which occur between the N-terminal PHD fingers and C-terminal RC region. Disruption of the N-C intramolecular interaction affects not only the size of PCLs phase separation particles but also the chromatin association of PRC2. Functionally, the CpG islands DNA can disrupt the N-C intramolecular interaction of PCLs to expose their middle IDRs, which in turn promote the PCLs driven PRC2 condensates formation, revealing a previously unrecognized role of PCLs in linking PRC2 and CpG islands chromatin via LLPS mechanism. This study provides a starting point for our further exploring how PRC2 accessory factors sense different chromatin environments, such as CpG islands or histone modifications, to regulate PRC2 mediated gene silencing via phase separation.

## Results

### PCLs undergo LLPS *in vitro* and *in vivo*

In the last decade, many types of “bodies” have been observed in the nucleus, such as PML bodies, Cajal bodies, and Polycomb or PcG bodies^30^. PcG bodies are large nuclear foci of PcG proteins that can be visualized under a microscope. However, to date, an ultrastructural description of the PcG bodies, such as their composition and functionality have not been completely understood. We wondered whether PRC2 core components or accessory factors can assemble within PcG bodies. To this end, we focused our initial attention on PRC2 accessory factors, PCL proteins (PCLs). We reconstitute *in vitro* droplet assays to answer whether PCLs can undergo LLPS. We previously determined that the C2 domain of SUZ12 (SUZ12-C2) was responsible for stable binding of the C-terminal RC region of PCLs^16^. In order to improve the stability of full-length PCLs, we first co-expressed and purified recombinant mEGFP-PHF19 with SUZ12-C2 binary complex in *Escherichia coli* (Supplementary Fig. 1A). When the sodium chloride concentration in the purified mEGFP-PHF19 with SUZ12-C2 binary complex solution was reduced from 1 M to 150 mM, micrometer-sized droplets arose that could be easily observed by fluorescence microscopy (Fig. 1A). The results indicate that high salt indeed prevented full-length PHF19 droplets formation *in vitro*.

**Fig. 1.**
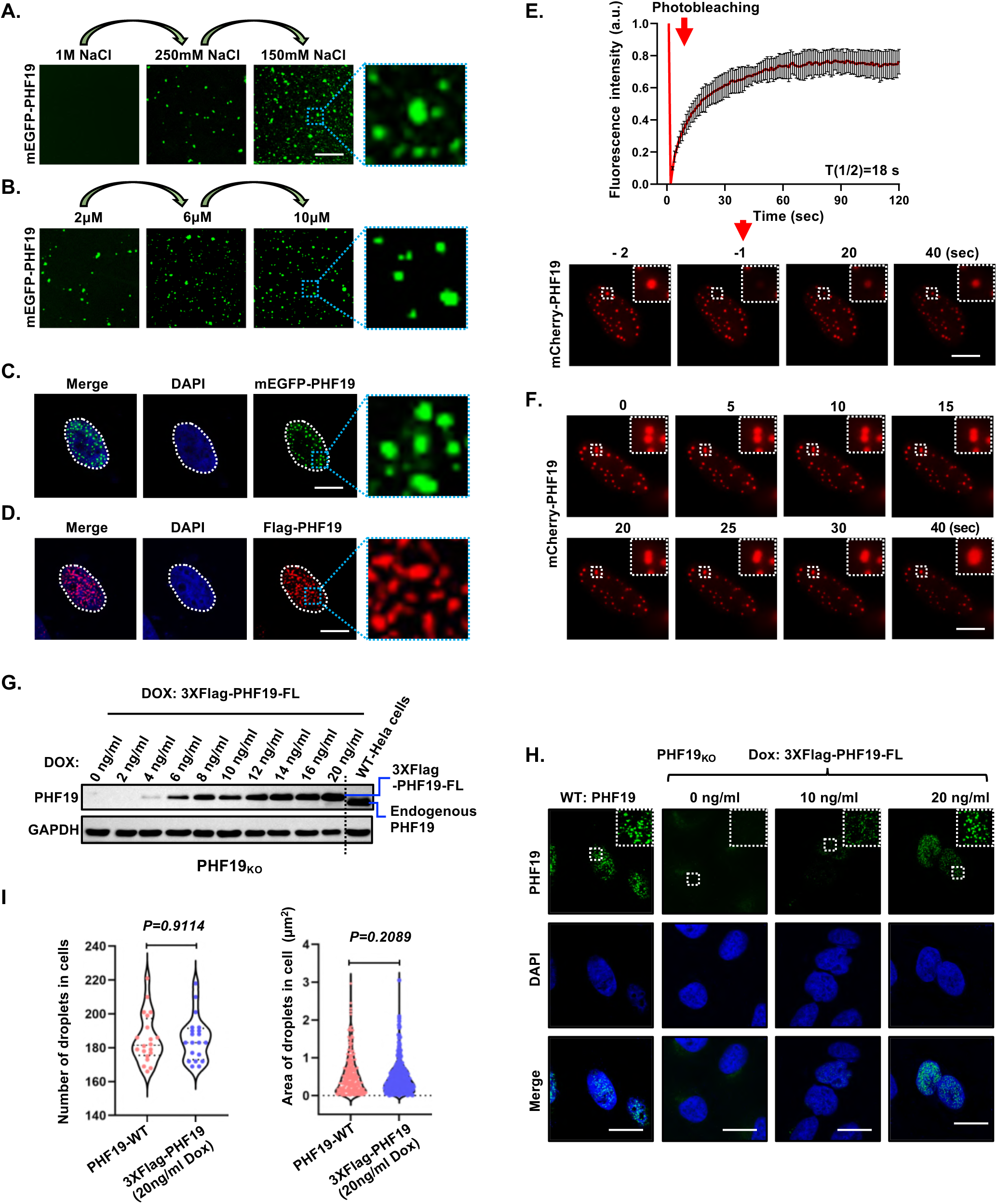
PHF19 phase separates into a liquid-like condensate *in vitro* and *in vivo*. **A** *In vitro* phase separation of purified mEGFP-PHF19-SUZ12(C2) binary protein complex (10 μM) at the indicated salt concentration. Scale bar,10 μm. **B** Representative images show that purified mEGFP-PHF19-SUZ12(C2) binary protein complex phase separation is dependent on protein concentration. Scale bars, 10 μm. **C** Fixed-cell images of mEGFP-tagged PHF19 in HeLa cells. Scale bars, 10 μm. **D** The Flag-PHF19 displays nuclear puncta in HeLa cells revealed by immunostaining with anti-Flag. Scale bars, 10 μm. **E** Immunostaining with anti-PHF19 antibody revealed that endogenous PHF19 in HeLa cells exhibited droplets. Scale bar, 10 μm. **F** FRAP analysis of the dynamics of mCherry-PHF19 droplets in HeLa cells. Representative images of fluorescence recovery are shown. Data were presented as mean values ± SEM (n = 3 independent experiments). Scale bar, 10 μm. **G** The fusion of mCherry-PHF19 droplets in HeLa cells is illustrated in live-cell images. Scale bar, 10 μm.

Similarly, the number and size of the droplets increased as the concentration of PHF19 was increased from 2 μM to 10 μM at 150 mM sodium chloride buffer condition (Fig. 1B). These phenomena are consistent with phase separation, which typically depends on salt and protein concentration. Thus, these results suggest that PHF19 can undergo phase separation to form condensates *in vitro*.

To test if PHF19 also forms biomolecular condensates in living cells, we ectopically expressed PHF19 protein by fused to monomeric EGFP (mEGFP) or Flag in HeLa cells. Both mEGFP-PHF19 and Flag-PHF19 form condensates in living cells (Fig. 1C and 1D), suggesting the formation of PHF19 condensates is irrespective of its carrying epitope tags. Next, we interrogate whether PHF19 condensates exhibit liquid-like property. We performed fluorescence recovery after photobleaching (FRAP) experiments on condensates of mCherry-PHF19 transient expressed in cells. FRAP data showed that 50% mCherry-PHF19 within condensates is recovered within 18 seconds (Fig. 1E), consistent with what are observed for other proteins that form either liquid-like condensates such as PRC1^7^ or membraneless organelles^31^. Furthermore, time-lapse confocal laser scanning showed these droplets could undergo fusion, these features are typically characteristic of liquid droplets (Fig. 1F).

To rule out the possibility that the PCLs condensates formation in Hela cells is due to their overexpression, we asked whether endogenous levels of PHF19 can form condensates. We applied CRISPR-mediated genome editing to knockout the endogenous PHF19 in HeLa cells (Supplementary Fig. 1C and 1D) and established a doxycycline (Dox)-regulated cell line with inducible, stable expression of 3XFlag- PHF19 gene in PHF19-KO HeLa cells. The results showed that the expression of 3XFlag-PHF19 protein level was gradually enhanced with increasing concentration of Dox. We determined the optimal level of Dox that induces expression of 3XFlag- PHF19 levels at near endogenous levels at 20 ng/mL Dox (Fig. 1G). We compared the condensates formation of PHF19 in these two cell lines (Wild-type Hela cells control and 20 ng/mL Dox induction of 3XFlag-PHF19 expressed in PHF19-KO Hela cells) and found no significant different condensates between these two cell lines (Fig. 1H and 1I). Together, these characteristics confirmed that PHF19 generates dynamic puncta by LLPS in living cells.

Similar to what was observed in PHF19, the formation of another two PCLs, PCL1/PHF1 and PCL2/MTF2 droplets *in vitro* also exhibit salt concentration dependence (Supplementary Fig. 1E). In addition, either ectopically expressed or endogenous PHF1 and MTF2 form a large number of condensates in living cells (Supplementary Fig. 1F and 1G). FRAP of PHF1 and MTF2 droplets show a fast recovery rate, indicating that PHF1 and MTF2 condensates are also highly dynamic and readily exchange with surrounding environment in living cells (Supplementary Fig. 1H and 1I). Taken together, the above results demonstrate that PCL proteins form biomolecular condensates and exhibit liquid-like properties *in vitro* and *vivo*.

#### PCLs form condensates colocalization with PRC2 and H3K27me3 *in vivo*

PCLs were known to interact with SUZ12 and exist in a stable complex with PRC2^21, 24, 25^. Hence, we investigated whether PCLs condensates colocalize with EZH2, a core component of PRC2 complex *in vivo*. To do this, we transfected HeLa cells with mCherry-PHF19 and stained endogenous EZH2 as well as mCherry-PHF19. Consistent with previous findings, we found that EZH2 formed condensates in living cells (Fig. 2A)^23^. The immunofluorescence imaging showed mCherry-PHF19 formed puncta that overlapped with EZH2 (Fig. 2A) and colocalized at H3K27me3, a chromatin mark produced by PRC2 complex (Fig. 2B), suggesting that PHF19 drives PRC2 complex condensates which are recruited to PcG-targeted genes or vice versa. By contrast, PHF19 forms far fewer puncta overlapped with H3K36me3-marked chromatin (Fig. 2C and 2D), which is basically consistent with previous reports that H3K27me3 and H3K36me3 only coexist ∼0.078% of H3 molecules^32^. Similarly, PHF1 and MTF2 form a large number of biomolecular condensates in the nucleus, which also colocalize with the EZH2 and H3K27me3 *in vivo* (Supplementary Fig. 2A). Thus, our results show that PCLs condensates colocalize with PRC2 complex and H3K27me3-marked chromatin regions *in vivo*.

**Fig. 2.**
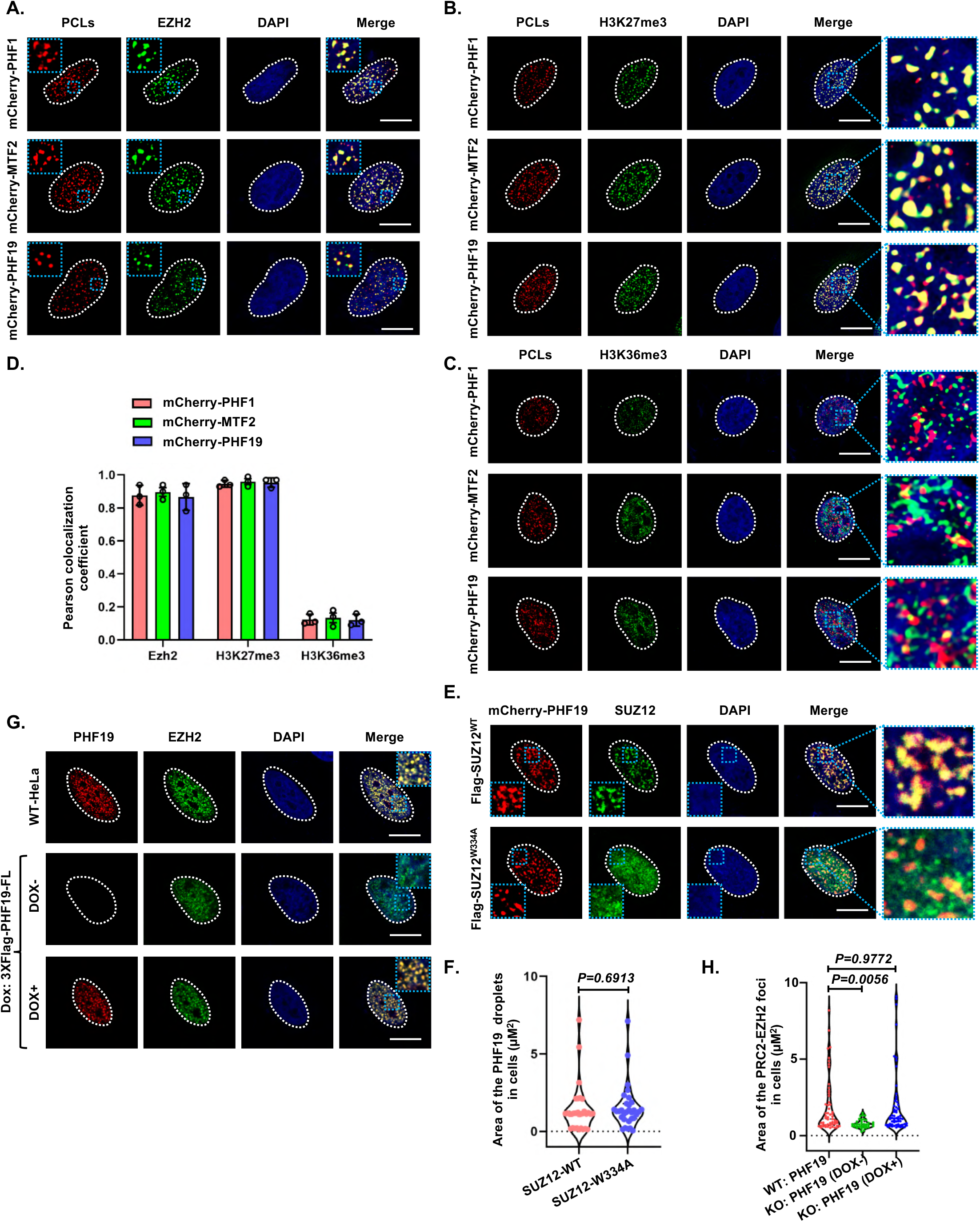
PCLs proteins form condensates colocalize with PRC2 and H3K27me3 *in vivo*. **A** Confocal images show colocalization of transfected mCherry-tagged PCLs with endogenous EZH2 in HeLa cells. Scale bars, 10 μm. **B** Confocal images illustrate colocalization of transfected mCherry-tagged PCLs with H3K27me3 in HeLa cells. Scale bars, 10 μm. **C** Representative images show that PCLs and H3K36me3 have low colocalization in HeLa cells. Scale bars, 10 μm. **D** Immunostaining show colocalization of mCherry-tagged PCLs with Flag-SUZ12^WT^, but not its mutants (Flag-SUZ12^W334A^) in HeLa cells. Scale bars, 10 μm.

Next, we asked whether the formation of PCLs nuclear puncta is dependent on PRC2 complex. Our previous crystal structure of PCL3/PHF19 containing PRC2 subcomplex demonstrated the W334A mutation of SUZ12 (SUZ12^W334A^) abolished the interaction of PHF19 with PRC2 complex^21^. To clarify the effect of PRC2 complex on PCLs condensates formation, we co-expressed PHF19 with wild-type (SUZ12^WT^) as well as mutant SUZ12 (SUZ12^W334A^) in HeLa cells. Compared with SUZ12^WT^, the formation of PHF19 condensates did not change in SUZ12^W334A^ co-transfected cells, indicative of the PCLs condensates formation independent on PRC2 complex (Fig. 2E and 2F). Conversely, whether the PCLs condensates formation can affect PRC2 phase separation in living cells. We compared the EZH2 condensates formation in the presence or absence of endogenous PHF19 in HeLa cells. The immunofluorescence assay results showed that knockout PHF19 reduced the degree of condensate formation of EZH2-PRC2 in HeLa cells, suggesting loss of PHF19 can inhibit the PRC2 condensates formation (Fig. 2G and 2H). Finally, we explore the possible correlation between PRC2 enzymatic activity and PHF19 condensates formation, we treated the mCherry-PHF19 expressing Hela cells with EPZ-6438, an EZH2 inhibitor. The results showed the cells treated with EPZ-6438 could lead to a global loss of H3K27me3 but did not change PHF19 droplets formation, suggesting PHF19 condensates formation is not relate to PRC2 enzymatic activity (Supplementary Fig. 2B-D). Thus, our results show that the formation of PCLs condensates are dispensable for PRC2 complex, but can drive PRC2 phase separation and colocalize on H3K27me3-dense chromatin.

#### The IDR domain of PCLs are required for their phase separation

Proteins that undergo LLPS often contain intrinsically disordered regions (IDRs) or low complexity domains (LCDs)^33, 34^. We used the prediction program PONDR- VSL2 and VS3 (http://www.pondr.com/) to analyze the sequence of PCLs. We observe a remarkably long region for PCL3/PHF19 with intrinsically disorder region (IDR) that contains 210 amino acids (residues from 320 to 530) and that was predicted to have a disorder score higher than 0.90 (Fig. 3A). Similarly, the IDRs were also observed in the middle region of PCL1/PHF1 and PCL2/MTF2 proteins (Supplementary Fig. 3A and 3B). To determine whether the IDRs of PCLs are required for their phase separation, we examined a series of truncated PCLs mutants with the aim of identifying the essential regions for driving PCLs condensates formation. We refer to the N-terminal Tudor domain and two adjacent plant homeodomain (PHD) fingers together with the extended homologous (EH) domain collectively as the TPE domain, the middle domain as the IDR domain, and C-terminal “reversed chromodomain’’ as RC region (Fig. 3B). Various mEGFP-PCLs constructs were generated and their expression were confirmed in Hela cells. We observed that the middle IDR of PHF19 formed larger droplets in the nucleus when compared with the N terminal TPE domain and the RC region of PHF19, respectively (Supplementary Fig. 3C). The similar phenomena were also observed in PHF1 and MTF2 (Supplementary Fig. 3D). These data support that the IDR domain is mainly involved in driving PCLs phase separation in living cells.

**Fig. 3.**
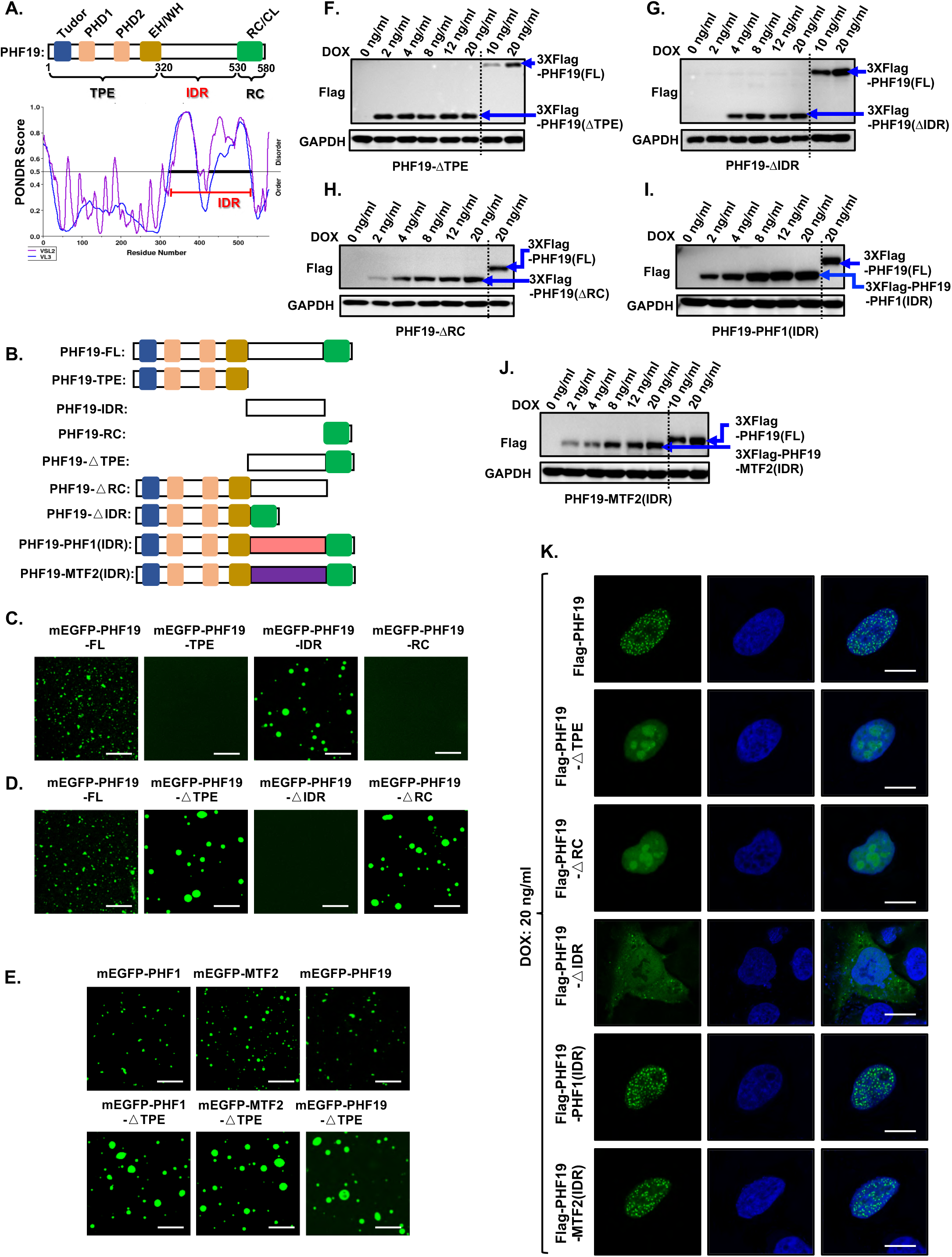
The IDR domain of PHF19 is required for condensate formation. **A** Diagram shows the structure of PHF19, including disordered regions predicted by the VSL2 and VL3 algorithm. IDR, intrinsically disordered region. **B** Schematic representation of PHF19 and mutants. TPE, Tudor domain and two adjacent plant-homeodomain (PHD) fingers together with the extended homologous (EH) domain. IDR, intrinsically disordered region. RC, the C-terminal “reversed chromodomain’’. **C** Representative images of PHF19 and its mutants droplets formation in HeLa cells. Scale bar, 5 μm. **D-E** *In vitro* phase separation of mEGFP-PHF19 and its mutants. Scale bar, 10 μm. **F** Representative images and quantitative analysis of the size of PHF19 formation phase separation droplets compared with its mutants. A two-tailed unpaired Student’s t-test was used for statistical analysis, and data are presented as mean values ± SEM (n= 50 independent samples). Scale bar, 5 μm. **G** *In vitro* phase separation of mEGFP-PHF1-SUZ12(C2), mEGFP-MTF2-SUZ12(C2), mEGFP-PHF19-SUZ12(C2) binary complexes and their corresponding mutants. Scale bar, 10 μm.

To further identify the direct contribution of the IDR domain to PHF19 phase separation, we constructed a panel of PHF19 deletion mutants fused to monomeric EGFP (mEGFP), expressed and purified them from *Escherichia coli*, and tested their condensates formation activity *in vitro* (Supplementary Fig. 1A and 1B). The mEGFP- PHF19-IDR formed numerous spherical droplets at 150 mM NaCl buffer condition (Fig. 3C). For comparison, the mEGFP-PHF19-TPE and mEGFP-PHF19-RC failed to form droplets under the same experimental conditions. Furthermore, deletion of the middle IDR region of PHF19 also fail to form droplets *in vitro* (Fig. 3D), further supporting that the IDR domain was critical for PHF19 condensates formation *in vitro*.

Interestingly, either deletion of the N terminal TPE domain or C terminal RC region of PHF19 promotes but not inhibits droplet formation as compared to the full-length PHF19 (Fig. 3D). Similar phenomenon was also observed *in vivo* (Supplementary Fig. 3E). To expand this observation for the whole PCL family proteins, we observed that either TPE or RC deletion of PHF1 and MTF2 formed larger particles *in vitro* and living cells (Fig. 3E and Supplementary Fig. 3F-3G). Thus, these results suggest that the middle IDR domain mainly contribute to PCLs driven phase-separated condensates formation, while the N and C terminus seem to be involved in controlling the size of PCLs condensate formation.

Again, to further demonstrate the IDR driven PCLs condensates formation are not due to their overexpression. We established Dox inducible cell lines with stable expression of 3XFlag tagged wild-type and a series of truncated PHF19 mutants in PHF19-KO Hela cells (Fig. 3B). We determined the optimal level of Dox that induces expression of wild-type and mutant PHF19 proteins at near endogenous levels at 20 ng/mL Dox (Fig. 1G and Fig. 3F-J). The immunofluorescence analysis showed either deletion of the N terminal TPE domain or C terminal RC region of PHF19 promotes droplet formation as compared to the full-length PHF19 at near endogenous levels (Fig. 3K). By contrast, loss of the middle IDR region of PHF19 was hardly to form puncta in nucleus (Fig. 3K). To emphasize the major role of IDR in driving PHF19 phase separation in living cell, we also replaced the IDR of PHF19 with the corresponding IDR from its homologs MTF2 or PHF1, respectively (Fig. 3B). The results showed that the condensates formation of the IDR deletion of PHF19 can be restored when re-adding with the IDRs from its homologs PHF1 or MTF2 (Fig. 3K), further suggesting the function of IDRs from PCL family protein are highly conserved, which are required for PCLs phase separation in living cells at near endogenous levels.

#### Intramolecular interaction regulates the phase separation of PCLs

Given that the N and C terminus need work together to control the proper PCLs condensates formation (Supplementary Fig. 3E, 3F and 3G). It is a reasonable hypothesis that the formation of PCLs condensate droplet size was dependent upon the presence of the N- and C- terminal domains together. To test the role of N and C terminus in controlling the droplet size of PCLs, we first used AlphaFold program to predict the structures of the full-length of PCLs. We found a large interaction interfaces between N-terminal TPE and C-terminal RC in PHF19 (Fig. 4A), as well as in PHF1 and MTF2 (Supplementary Fig. 4A and 4B). To validate this prediction, we first performed co-IP assays followed by immunoblot analyses in the Hela cells. The results showed that C-terminal RC region is able to interact with N-terminal TPE domain, but not with the middle IDR region (Fig. 4B). In agreement with the Co-IP data, the immunofluorescence analysis showed that PHF19-RC colocalizes with PHF19-TPE in cytoplasm, but not with PHF19-IDR in nucleus (Fig. 4C). Further mapping analysis showed either deletion of Tudor or EH domain does not affect the N and C terminus interaction and colocalization, suggesting PHF19-RC mainly interacts with the N-terminal PHD fingers (Fig. 4D and 4E). To further investigate whether the N- and C-terminus interaction is direct, glutathione S-transferase (GST)-tagged PHF1-RC and Flag-tagged PHF1-TPE were co-expressed in *Escherichia coli* for GST pull-down assays. The Coomassie blue staining clearly showed The PHF1-RC readily pulled down PHF1-TPE, suggesting that PHF1-RC directly interacts with PHF1-TPE (Supplementary Fig. 4C). Thus, these results together indicate that PCLs have intramolecular interactions mainly through the N-terminal PHD fingers and C-terminal RC region.

**Fig. 4.**
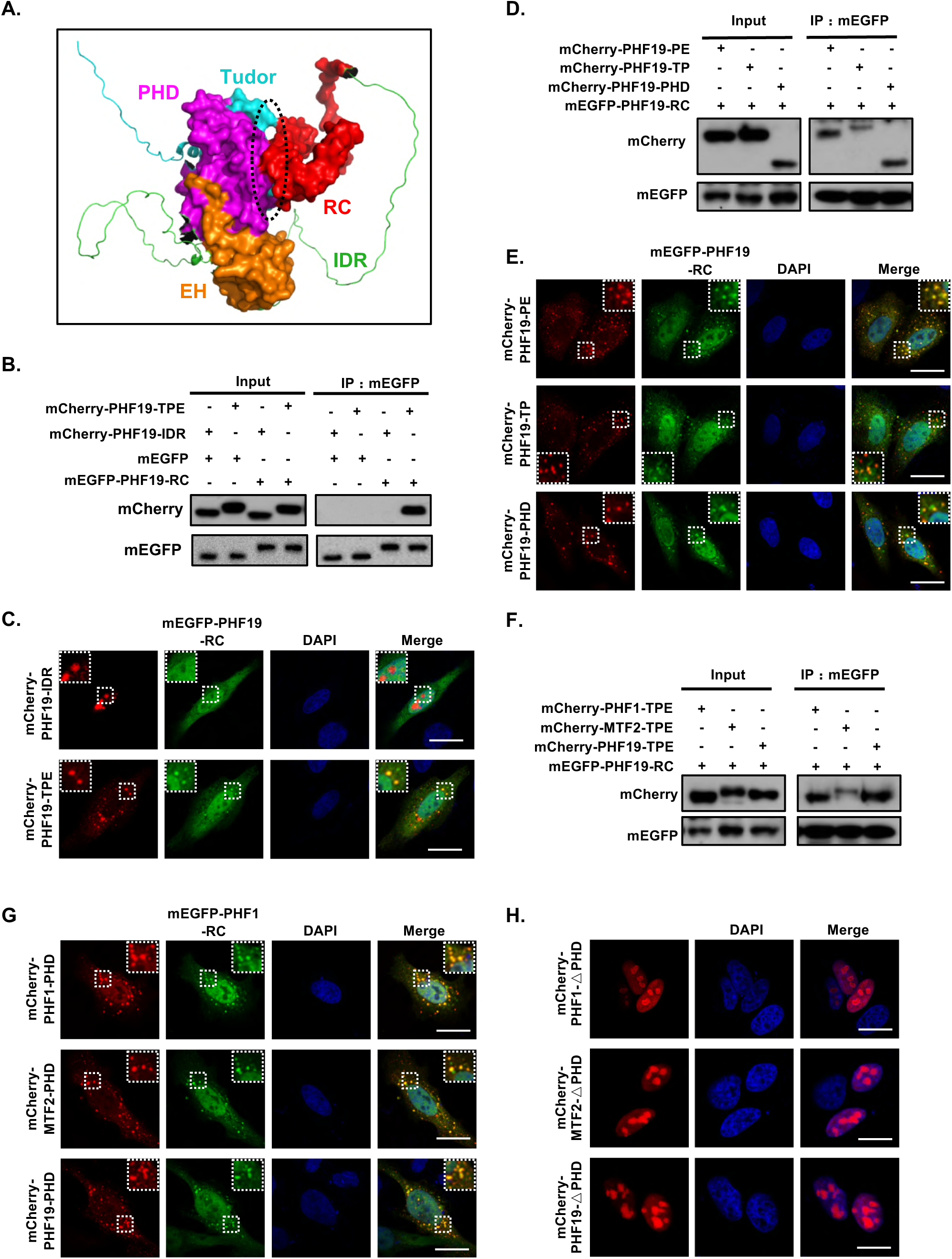
I**n**tramolecular **interaction regulates the phase separation of PCLs proteins. A** The structure of PHF19 protein was predicted by using the AlphaFold algorithm. The black dotted line indicates the protein interaction interface. **B** The interaction of mEGFP-PHF19-RC with mCherry-PHF19-TPE and mCherry- PHF19-IDR were analyzed in HeLa cells by co-immunoprecipitation (co-IP) assay. **C** Representative images show colocalization of transfected mEGFP-PHF19-RC with mCherry-PHF19-TPE or mCherry-PHF19-IDR in HeLa cells. Scale bars, 5 μm. **D** Co-immunoprecipitation (co-IP) was used to analyze mEGFP-PHF19-RC interactions with mCherry-PHF19-PE, mCherry-PHF19-TP, and mCherry-PHF19- PHD in HeLa cells. **E** Representative images show colocalization of transfected mEGFP-PHF19-RC with mCherry-PHF19-PE, mCherry-PHF19-TP, or mCherry-PHF19-PHD in HeLa cells. Scale bars, 5 μm. **F** The mEGFP-PHF19-RC interaction with mCherry-PHF1-TPE, mCherry-MTF2-TPE and mCherry-PHF19-TPE were analyzed in HeLa cells by co-immunoprecipitation (co-IP) assay. **G** Representative images show colocalization of transfected mEGFP-PHF1-RC with mCherry-PHF1-PHD, mCherry-MTF2-PHD, or mCherry-PHF19-PHD in HeLa cells. Scale bars, 5 μm. **H** Representative image show droplets formation of mCherry-PHF1(△PHD), mCherry- MTF2(△PHD) and mCherry-PHF19(△PHD). Scale bars, 5 μm

Next, we ask whether there are heterologous interactions between the N terminus from one PCL protein and C terminus from another PCL protein, we co-expressed mEGFP-PHF19-RC with mCherry-PHF1-TPE, mCherry-MTF2-TPE or mCherry- PHF19-TPE (as control) in Hela cells, respectively. The co-immunoprecipitation (Co-IP) followed by western blot assays showed that PHF19-RC could interact with PHF1- TPE or MTF2-TPE (Fig. 4F), just as it interacts with PHF19-TPE control. The immunofluorescence assay showed PHF19-RC colocalized with PHF1-TPE or MTF2- TPE droplets in the cytoplasm as well as with PFH19-TPE (Supplementary Fig. 4D). The same phenomena were also observed in PCL1/PHF1 and PCL2/MTF2 proteins (Fig. 4G and Supplementary Fig. 4D). Actually, the heterologous interactions from different PCL proteins were also observed in the context of full-length PCLs (Supplementary Fig. 4F). Finally, we sought to demonstrate whether the N-C intramolecular interaction of PCLs control their condensate droplets size, we constructed PCLs mutant with PHD fingers deletion, which disrupt the N-C intramolecular interaction of PCLs. Immunofluorescence assays showed that deletion of the PHD fingers resulted in the formation of larger droplets of PCLs (Fig. 4H), similar as observed in TPE deletion of PCLs mutants (Supplementary Fig. 3G and 3H). Likewise, overexpression of PHF19-RC to disrupt the PHF19 intramolecular interaction also leads to form larger particles of PHF19 protein condensates. (Supplementary Fig. 4G-H). Together, these results indicated that disruption of the intramolecular interaction promote the PCLs phase separation.

PCLs utilize their C-terminal RC region to interact with SUZ12-C2 domain and incorporate into PRC2 complex^16, 21^. We ask whether the binding of SUZ12 to PCL proteins affect the N-C intramolecular interaction of PCLs in the context of PRC2-PCLs complexes. To answer this, we used anti-GFP nanobody coupled beads to capture mEGFP-PHF19-RC protein and examined the binding of Flag-PHF19-TPE protein to mEGFP-PHF19-RC protein with increasing amounts of purified SUZ12-C2 protein. We observed that SUZ12-C2 did not disrupt the interaction between TPE and RC *in vitro* (Fig. 5A), indicating the N-C intramolecular interaction of PCLs should be reserved in PRC2-PCLs complexes. The same conclusion was also obtained from the *in vitro* GST-PHF1-RC pull down PHF1-TPE with or without the presence of SUZ12- C2 protein (Supplementary Fig. 4C). In addition, the droplet size of PHF19 phase separation does not change when PHF19 co-expressed with SUZ12^WT^ as well as SUZ12^W334A^, which SUZ12 mutant was demonstrated to lose the ability to bind to PHF19^21^ (Supplementary Fig. 5A-B). Together, these results suggest that the N-C intramolecular interaction of PCLs should be still reserved when PCL proteins incorporate into PRC2 complex. We failed to resolve the crystal structure of the N-terminal TPE (PHF1-TPE) bound to C-terminal RC region (PHF1-RC) complex. In future, high resolution cryo-EM structure of full-length PCLs containing PRC2 complex will help to answer how PCLs N terminal TPE domain interact with the C terminal RC region in the context of PRC2 complex in more detail.

**Fig. 5.**
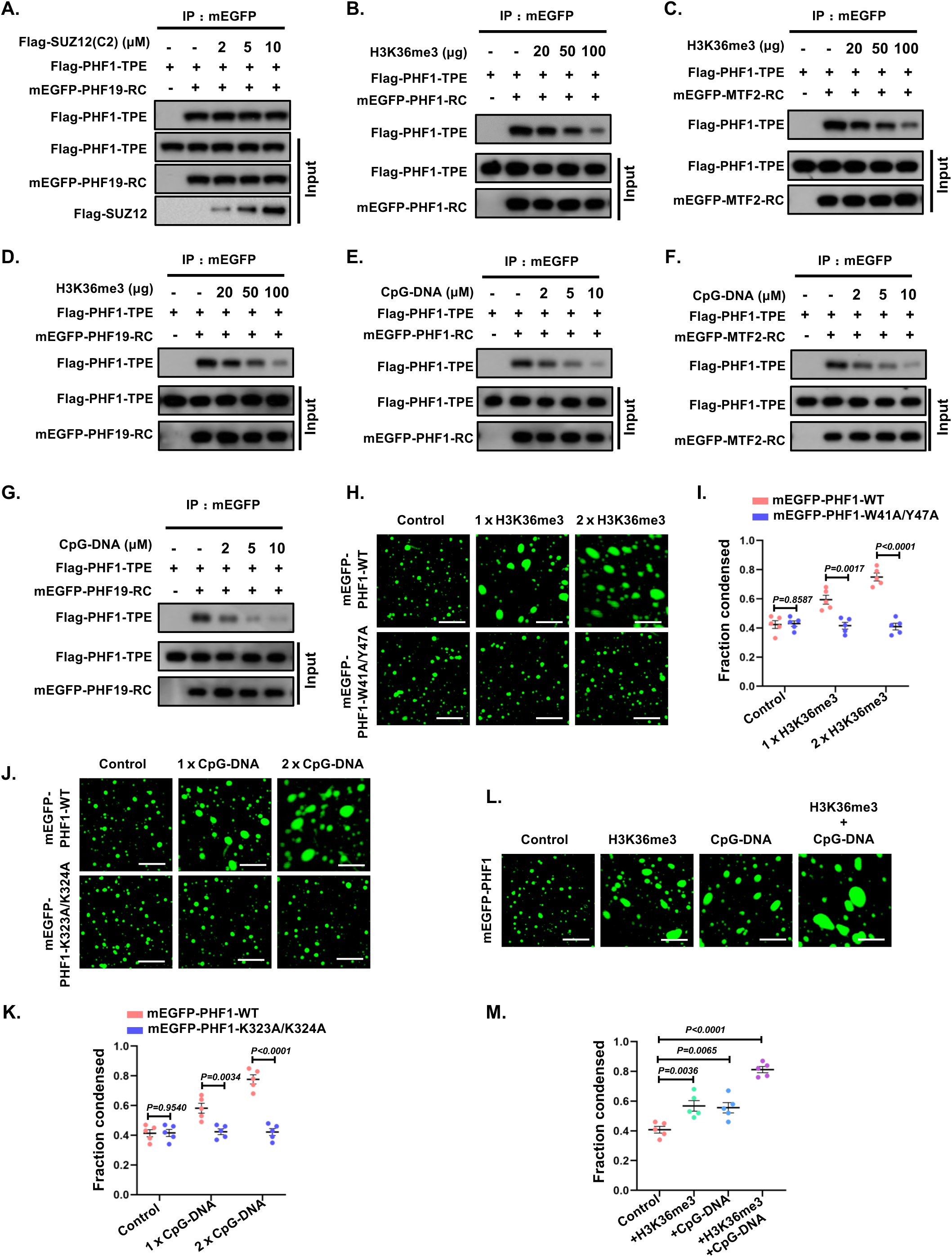
Effect of H3K36me3 **and CpG-DNA on phase separation of PCLs proteins. A** Effect of Flag-SUZ12(C2) on the interaction between Flag-PHF1-TPE and mEGFP- PHF19-RC *in vitro*. **B-D** Effect of H3K36me3 peptide on the interaction between Flag-PHF1-TPE and mEGFP-PHF1-RC, mEGFP-MTF2-RC, mEGFP-PHF19-RC *in vitro*. **E-G** Effect of CpG-DNA on the interaction between Flag-PHF1-TPE and mEGFP- PHF1-RC, mEGFP-MTF2-RC, mEGFP-PHF19-RC *in vitro*. **H-I** Representative images and quantitative analysis show that H3K36me3 peptide regulated the phase separation of mEGFP-PHF1 and mEGFP-PHF1^W41A/Y47A^ mutants *in vitro*. The one-way ANOVA was used for statistical analysis, and data are presented as mean values ± SEM (n= 50 independent samples). Scale bar, 10 μm. **J-K** Representative images and quantitative analysis show that CpG-DNA promoted mEGFP-PHF1 phase separation *in vitro*, but not mEGFP-PHF1^K323A/K324A^ mutants. The one-way ANOVA was used for statistical analysis, and data are presented as mean values ± SEM (n= 50 independent samples). Scale bar, 10 μm. **L-M** Representative images and quantitative analysis showed the effect of H3K36me3 and CpG-DNA on mEGFP-PHF1 phase separation *in vitro*. The one-way ANOVA was used for statistical analysis, and data are presented as mean values ± SEM (n= 50 independent samples). Scale bar, 10 μm.

#### CpG islands DNA trigger PCLs phase separation via disruption of intramolecular interaction

The apo-form structure of the PHF1-TPE show that the Tudor, PHD fingers, and EH domain organize into a compact structural architecture^26^. This raises the question of whether Tudor domain binding to H3K36me3 modified histone peptide or EH domain binding to CpG-DNAs disrupts the N-C intramolecular interaction of PCLs and promotes their phase separation. To do this, we used mEGFP nanoantibody coupled beads to capture mEGFP-PHF1-RC and then examined the binding of Flag-PHF1-TPE to mEGFP-PHF1-RC in the presence of H3K36me3 histone peptide or CpG-DNAs, respectively. Our pull-down results clearly showed that PHF1-RC can directly interact with PHF1-TPE, while H3K36me3 histone peptide and CpG-DNAs inhibit their interaction in a concentration dependent manner (Fig. 5B and 5E). Similarly, the heterologous interactions of PCL proteins were also inhibited by the H3K36me3 histone peptide or CpG-DNAs in a concentration-dependent manner (Fig. 5C-D and 5F-G). Inspired by the crystal structure of the PHF1-TPE bound CpG-DNAs in the presence of H3K36me3 histone peptides^26^, we further explore whether H3K36me3 peptide and CpG-DNAs have a coordinated effect on disruption of the intramolecular interaction of PHF1 protein. The pull-down results clearly show H3K36me3 peptide and CpG-DNAs synergize to disrupt the N-C intramolecular interaction of PHF1 (Supplementary Fig. 5C and 5D).

Next, we wondered whether H3K36me3 peptide or CpG-DNAs can regulate PCLs condensate formation and size via disruption of their intramolecular interaction. The W41A or Y47A mutation of PHF1 was previously demonstrated to inhibit the binding of H3K36me3 peptides to the Tudor domain^35^, while K323A or K324A mutation of PHF1 inhibit CpG-DNAs binding to the EH domain^26^. As control experiments, we co-expressed and purified these two mutant proteins (referred to as PHF1^W41A/Y47A^ and PHF1^K323A/K324A^) with SUZ12-C2 respectively in *Escherichia coli* and then performed phase separation assays *in vitro*. Compared with the wild-type PHF1 (PHF1^WT^), these two mutants PHF1^W41A/Y47A^ and PHF1^K323A/K324A^ did not cause any significant changes in the size of PHF1 condensates (Supplementary Fig. 5E), implying that these amino acid mutations did not affect PHF1 phase separation in the absence of H3K36me3 peptides or CpG-DNAs. However, when adding H3K36me3 peptides, we found that the wild-type PHF1 protein formed larger condensates when compared with PHF1^W41A/Y47A^ protein (Fig. 5H, 5I and Supplementary Fig. 5F). Likewise, CpG-DNA promotes PHF1^WT^ but not PHF1^K323A/K324A^ condensates formation (Fig. 5J, 5K and Supplementary Fig. 5F). Consistent with the *in vitro* droplet assays, immunofluorescence results showed that wild-type PHF1 formed larger droplets in cells than PHF1^W41A/Y47A^ and PHF1^K323A/K324A^ mutants, and had significant association with chromatin, suggesting change PHF1 phase separation affect its chromatin association (Supplementary Fig. 5G and 5H). To further investigate whether H3K36me3 levels affect PHF19 phase separation *in vivo*, we overexpressed a demethylase NO66 and its dead mutant (NO66^H340A/H405A^) with PHF19 in cells. We observed that overexpression of NO66 reduced the droplet size of PHF19 phase separation (Supplementary Fig. 5I and 5J), but not the NO66^H340A/H405A^ mutants, indicating that reduced H3K36me3 levels attenuated the inhibitory effect on the N-C intramolecular interaction of PHF19, thereby decreasing PHF19 phase separation droplets size. Also, H3K36me3 peptides acts in synergy with CpG-DNAs to promote PHF1 condensates formation, consistent with observations that they have a coordinated effect on disrupting the PHF1 intramolecular interaction (Fig. 5L and 5M). Thus, these results support that the CpG-DNAs or H3K36me3 peptides promotes the phase separation of PCLs likely via disrupting their N-C intramolecular interaction.

#### CpG islands triggers the condensation of PRC2 via PCLs

PCLs recognize CpG islands DNA through their N-terminal EH domain and guide PRC2 towards specific genomic loci for gene silencing^26^. Our in vitro phase separation experiments showed that PRC2 could be readily trapped into the PHF1 driven phase separation droplets (Fig. 6A and 6B), suggesting the phase separation of PHF1 facilitates the formation of PRC2 condensates. Given that CpG-DNAs can promote the phase separation of PCLs via disruption of their intramolecular interaction, we now ask whether the CpG-DNAs can trigger PRC2 condensates formation via PCLs. To answer this question, we first explore the effect of the intensity of CpG islands DNA on the formation of PCLs phase separation. We use different lengths of a natural mouse CpG islands DNA from the LHX6 gene that was used and described as previously^21^, including 153bp, 353bp and 799bp (Fig. 6C). The results show that CpG-DNAs promote PHF1 phase separation depending on DNA length (Fig. 6C-D), suggesting the intensity of CpG islands DNA regulates the degree of PCLs condensates formation likely via multivalent interactions between PCLs and CpG-DNAs (Supplementary Fig. 6A-B). Similarly, MTF2 and PHF19 can also trap CpG-DNAs into their driven phase separation droplets (Fig. 6E). Next, we investigate whether the CpG-DNAs trigger the PRC2 condensates formation through PCLs phase separation, we performed the *in vitro* phase separation experiments with the different combination of PRC2, CpG-DNAs and PHF1. It was observed that PRC2 complex did not form droplets when incubated with CpG-DNAs only, while the addition of PHF1 significantly promoted condensates formation from PRC2 (Fig. 6F). This result shows that the formation of PRC2 condensates on CpG islands is highly dependent on PHF1 protein. Indeed, the full-length, but not the N terminal TPE domain deletion of PHF1 recruit CpG-DNAs into their driven phase separation droplets (Supplementary Fig. 6C and 6D), suggesting that CpG-DNA promotes PHF1 phase separation via the N terminal TPE domain of PCLs binding to CpG-DNA. In addition, we also observed that CpG-DNAs promote the phase separation of PRC2-PHF1 complex when compared with PRC2 complex (Fig. 6G-H), further supporting of CpG islands DNA triggers the PRC2 condensates formation dependent upon PCLs.

**Fig. 6.**
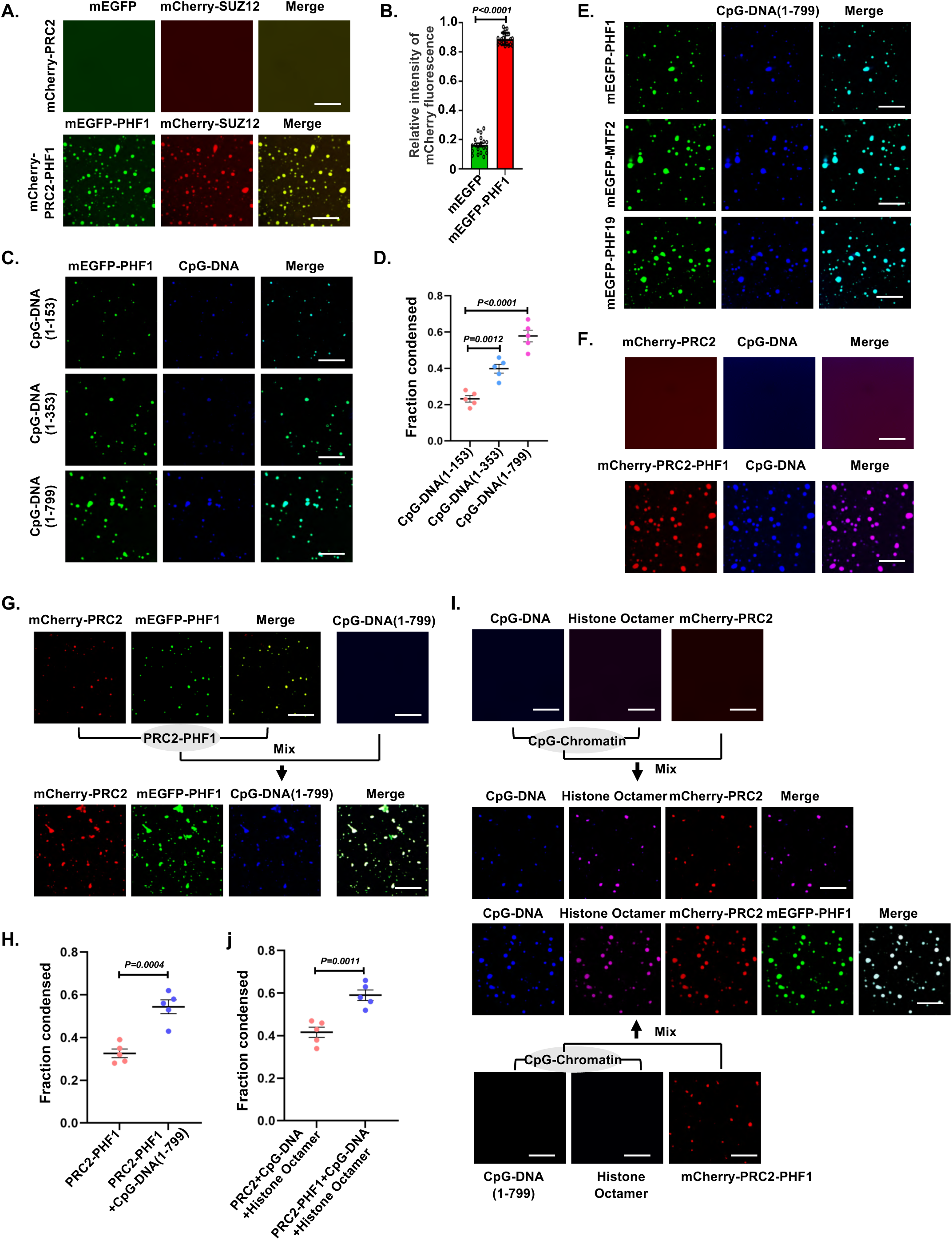
PHF1 phase separation promotes the condensation of PRC2 to CpG-DNA chromatin. **A** Representative images show that PHF1 phase separation recruits PRC2 into condensation. The SUZ12 is a core component of PRC2 complex labeled with mCherry tag. Scale bars, 10 μm. **B** Quantitative analysis of PHF1 phase separation recruits PRC2 into condensation. A two-tailed unpaired Student’s t-test was used for statistical analysis, and data are presented as mean values ± SEM (n= 20 independent samples). **C-D** Representative images and quantitative analysis showed the effect of CpG-DNA length on mEGFP-PHF1-SUZ12(C2) binary protein complex phase separation. CpG-DNA amplification primer was labeled with pacific blue. A two-tailed unpaired Student’s t-test was used for statistical analysis, and data are presented as mean values ± SEM (n= 50 independent samples). Scale bars, 10 μm. **E** Representative images show that mEGFP-PHF1-SUZ12(C2), mEGFP-PHF19- SUZ12(C2), and mEGFP-MTF2-SUZ12(C2) proteins phase separation recruits CpG-DNA into condensation. Scale bars, 10 μm. **F** Representative images showed that PRC2 is recruited to PHF1 phase separation condensates. Scale bars, 10 μm. **G-H** Representative images and quantitative analysis show that CpG-DNA promotes phase separation of PRC2-PHF1 complex. A two-tailed unpaired Student’s t-test was used for statistical analysis, and data are presented as mean values ± SEM (n= 50 independent samples). Scale bars, 10 μm. **I-J** Representative images and quantitative analysis show that PHF1 phase separation promotes the localization of PRC2 to CpG-DNA chromatin. Histone octamer are labeled with Cy5 dye. A two-tailed unpaired Student’s t-test was used for statistical analysis, and data are presented as mean values ± SEM (n= 50 independent samples). Scale bars, 10 μm.

In mammalian, the CpG islands were often associated with nucleosome-free promoters^36^. Here, we propose that PCLs drive PRC2 condensates formation on the CpG islands DNA, which compacts and brings two adjacent nucleosomes closer together to enhance the nucleosome density and facilitate PRC2 depositing the H3K27me3 on corresponding CpG islands chromatin. To test this possibility, we reconstitute the CpG islands chromatin by mixing the CpG islands containing DNA (1- 799bp) and histones octamer *in vitro*. Compared with PRC2 complex alone, PRC2- PHF1 formed larger droplets in the presence of CpG islands chromatin (Fig. 6I and J). These results further support that PHF1 promoted the formation of PRC2 condensates on CpG islands chromatin by phase separation *in vitro*.

#### PCLs Phase separation affects PRC2 recruitment and catalytic activity

PCLs either modulate PRC2 catalytic activity or recruit PRC2 to its genomic targets partially by recognizing CpG islands chromatin. We last sought to determine whether the phase-separated PCLs condensates formation can function to regulate PRC2 enzymatic activity or its chromatin association. Firstly, the immunofluorescence assay of the full-length PCL2/MTF2 showed punctate nuclear foci overlapping with DAPI-dense regions of chromatin (Fig. 7A and 7B). While the N or C terminus deletion of MTF2 mutants showed larger droplets occurred at regions with low DNA density, implying the aberrant of MTF2 phase-separated condensates formation will result in its chromatin dissociation (Fig. 7A and 7B). Actually, similar phenomena were previously observed for various IDRs containing nuclear proteins, which phase separate into larger biomolecular condensates that mechanically exclude chromatin^37^. Additionally, the N terminal TPE deletion of MTF2 also switches the localization of EZH2 from DAPI-dense regions of chromatin to regions of low chromatin density (Fig. 7A and 7B), indicating PRC2 chromatin binding is closely correlated with the degree of MTF2 phase separation. To further identify the role of MTF2 phase-separation in PRC2 chromatin association, we expressed the PHD fingers deletion of MTF2 mutant which loss of the N-C intramolecular interaction of PCLs and analyzed PRC2 chromatin association in Hela cell. Immunofluorescence assays showed deletion of PHDs result in larger MTF2 condensates formation, concomitant with loss of H3K27me3 colocalization and chromatin association (Fig. 7C and 7D). Moreover, the nuclear localization of EZH2 was also found to switch into low chromatin density regions, further supporting change of the MTF2 phase separation affect PRC2 chromatin association (Fig. 7A and 7B). Similar dynamic behavior for PHF1 and PHF19 mutants are also observed (Supplementary Fig. 7A-D). To demonstrate whether the N or C terminus deletion of PCLs mutants mislocalize to the nucleolar, we performed immunofluorescence assay to detect the colocalization of wild-type MTF2 or its truncated mutants with nucleolar marker NPM1. The results showed that NPM1 colocalizes well with all truncated MTF2 mutants in nucleolar, suggesting the aberrant condensates formation of MTF2 mutants lead to MTF2 mislocalization to the nucleolar (Supplementary Fig. 7E). The similar immunofluorescence phenomenon was observed in PHF19 protein (Supplementary Fig. 7F). These observations implicated the chromatin association of PRC2 to is closely correlated with the degree of PCLs phase-separated states.

**Fig. 7.**
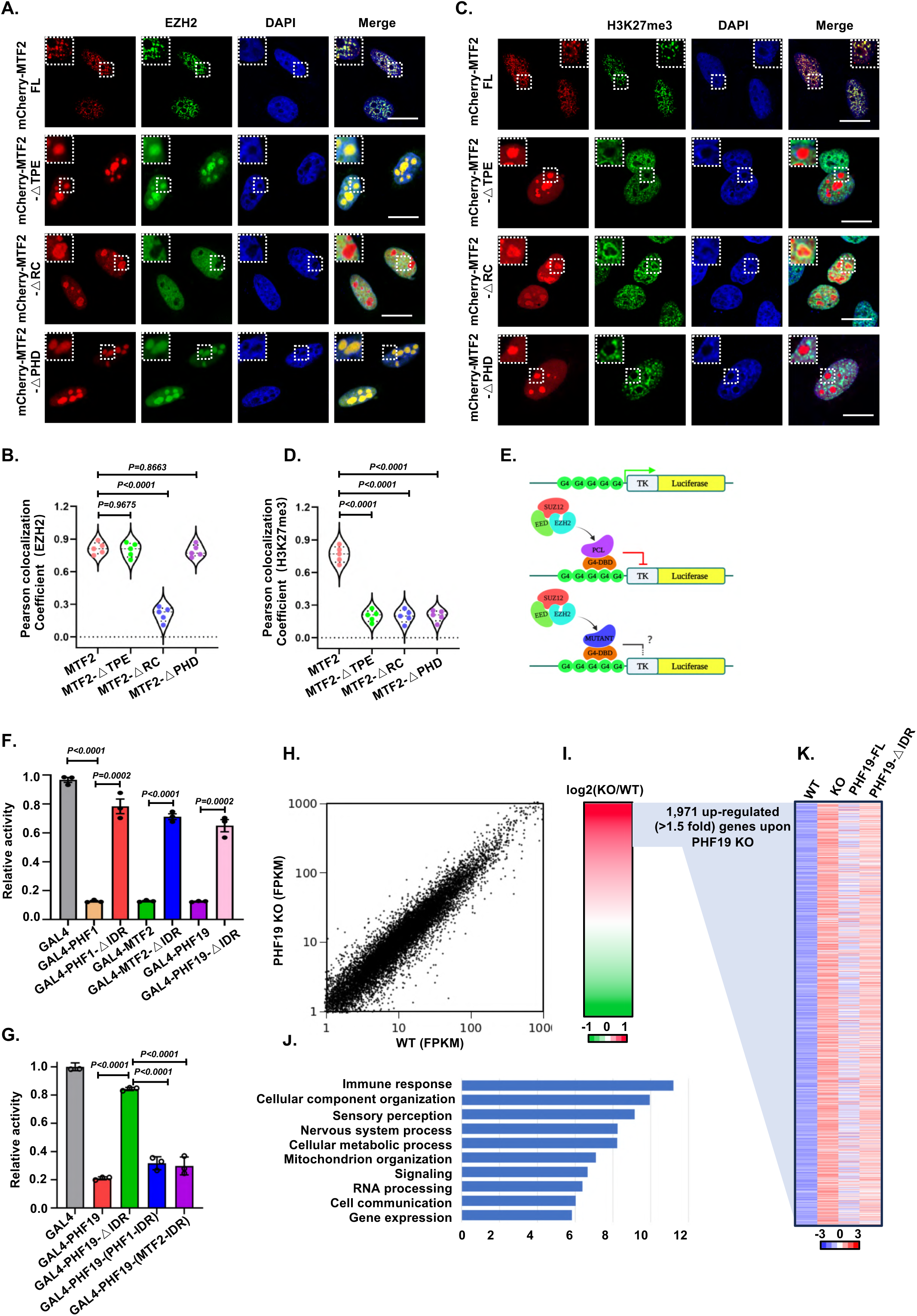
PCLs phase separation regulates PRC2 localization chromatin and transcriptional repression. **A** Representative images show colocalization of transfected mCherry-MTF2 and its mutants with endogenous EZH2 and chromatin marked by DAPI in HeLa cells. Scale bars, 5 μm. **B** Representative images show colocalization of transfected mCherry-MTF2 and its mutants with H3K27me3 in HeLa cells. Scale bars, 5 μm. **C** Schematic of PCLs and its mutants proteins based on GAL4 reporter gene repression assay. **D** Luciferase gene repression of PHF19 and its mutants. The one-way ANOVA was used for statistical analysis, and data are presented as mean values ± SEM (n= 3 independent experiments). **E** Effects of IDR region on the PCL-mediated gene repression were assayed. The one-way ANOVA was used for statistical analysis, and data are presented as mean values ± SEM (n= 3 independent experiments). **F** Effects of IDR region on the PHF19-mediated gene repression were assayed. The IDR of PHF19 were replaced by those of PHF1-IDR and MTF2-IDR, respectively. The one-way ANOVA was used for statistical analysis, and data are presented as mean values ± SEM (n= 3 independent experiments).

Next, we ask whether the PCLs phase separation can regulate PRC2 catalytic activity. As a starting point, we generated human GAL4-PCLs fusion proteins and their mutants in a HEK293 cell line that contained an integrated heterologous GAL4-luciferase reporter gene under control of a “5×GAL4UAS-Thymidine Kinase (TK) promoter” cassette (Fig. 7E)^28, 38^. We observed that PHF1, MTF2 and PHF19 were able to recruit PRC2 to suppress the reporter gene through their C-terminal RC regions (Supplementary Fig. 7G-I), as removal of the C-terminal RC region abolished gene repression. Compared with full-length PHF19, the deletion of the IDR domain reduced the extent of reporter gene suppression (Fig. 7F), suggesting the IDR driven PHF19 phase separation involve in regulating PRC2 catalytic activity. The similar results were observed for PHF1 and MTF2 mutants (Fig. 7F). In order to further emphasize the role of IDR-driven PCLs phase separation in regulating PRC2 catalytic activity, we replaced the IDR of PHF19 with the corresponding IDR from its homologs MTF2 or PHF1, respectively. Interestingly, the results showed that the inhibition effect of the IDR deletion of PHF19 on reporter gene can be restored when re-adding with the IDRs from its homologs PHF1 or MTF2 (Fig. 7G), suggesting the function of IDRs from PCL family protein are highly conserved, which are involved in regulating PRC2 enzymatic activity via phase separation. Finally, we performed transcriptome profiling to evaluate the role of PHF19-IDR-dependent phase separation on PRC2 target genes regulation. Our RNA-Seq analysis identified 1971 up-regulated differently expressed genes (DEGs) in PHF19-KO Hela cells (Fig. 7H-J). Rescued PHF19-KO Hela cells with a 3XFlag- PHF19-FL, but not 3XFlag-PHF19-ΔIDR mutant can re-silence these up-regulated expressed genes observed in PHF19-KO Hela cells. (Fig. 7J). Thus, our whole transcriptome sequencing data strongly indicated the PHF19 regulated PRC2 target genes silencing is highly dependent on its IDR region. Gene ontology (GO) analysis was further used to show that upregulated genes are involved in essential biological processes, such as immune response, nervous system processing, various signaling pathways, and cellular component organization (Fig. 7K). These observations indicated the PHF19-IDR-dependent phase separation regulates the PRC2 target genes expression.

## Discussion

Growing evidence has demonstrated that phase separation plays a critical role in gene regulation^39, 40^, a process by which molecules, such as proteins and nucleic acids, are concentrated in membrane-less compartments. Phase separation has long been implicated in PRC1 mediated gene silencing, which it was proposed to utilize the LLPS to compact chromatin and silence gene expression^6, 7, 8, 9^. PRC1 and PRC2 usually co-occupy thousands of shared target genes and cooperate to repress genes expression. In this regard, we first identified the PRC2 accessory factor, PCLs contain the previously unidentified IDR domains, which drive the formation of PCLs biomolecular condensates. Compared with highly-ordered structures, IDRs lacks stable structural features^41^, and thus has a high degree of flexibility to interact with other proteins to form condensates. Here, our biochemical data show PCLs control the condensate size via their intramolecular interaction, which occur between the N-terminal PHD fingers and C-terminal RC region. The N-C intramolecular interaction of PCLs likely limit the flexibility of the middle IDR region and thus inhibit its ability to form large droplets. As we observed, the loss of N terminal TPE domain or C terminal RC region, resulted in enhanced PCLs droplet size. Furthermore, deletion of the PHD finger to disrupt the N-C intramolecular interaction of PCLs also promote PCLs condensates formation. Thus, we first showed that PCLs phase-separated biomolecular condensates formation can be regulated through their intramolecular interactions, in contrast to several previous studies which have focused on post-translational modifications or mutations of target proteins to change proteins phase separation ^42, 43^.

PCLs play a critical role in guiding PRC2-catalyzed H3K27me3 on specific genomic loci by recognizing H3K36me3 histone mark or CpG islands DNA through Tudor and EH domain, respectively^24, 26^. How PRC2 complex access active chromatin to lead to *de novo* gene silencing during lineage transitions are still not understood completely. Previous studies proposed PCLs recruit the demethylase NO66 in transcriptionally active chromatin to de-methylate the H3K36me3 and re-silence the active transcribed genes expression via their N terminal Tudor recognizing the H3K36me3^27, 28^. Meanwhile, the H3K36me3 was also found to inhibit the enzymatic activity of PRC2 to prevent PRC2 over-repress the active transcribed genes expression^44, 45, 46^. The cryo-EM structure of PRC2-PHF1 bound H3K36me3 modified dinucleosome reveals the binding of EZH2 to nucleosomal DNA orients the H3 tail and threads H3K27 into its active site, in which the H3K36 occupies a critical position in the EZH2- nucleosomal DNA interface. The methylated H3K36 (H3K36me3) affect the above interactions between EZH2, the nucleosomal DNA and the H3 tail therefore permits allosteric inhibition of PRC2 in transcriptionally active chromatin^46^. Remarkably, the enzymatic activity of PRC2 in transcriptionally active chromatin should be tightly controlled. In this study, we further showed the recognition of H3K36me3 by Tudor domain disrupts the N-C intramolecular interaction to facilitate the PCLs driven PRC2 phase-separated formation, which add an additional layer of complexity in PRC2-PCLs complexes mediated *de novo* gene silencing. Thus, these results presented here suggest that H3K36me3 inhibits PRC2 enzymatic activity not only via affecting the interaction between EZH2 CXC domain and nucleosomal DNA, but also through regulating condensation behaviors of PRC2 on chromatin. This study establishes a pioneering example in which chromatin environment can trigger PCLs driven PRC2 condensation behaviors formation and offers a model for our understanding the role of different chromatin environment in PRC2 functional regulation via changing its accessory factor condensate formation. Detailed investigations into the aforementioned intramolecular interactions will further illuminate how PCLs condensates form and inform strategies to perturb these condensates.

PCLs have been shown to recognize CpG islands DNA and link PRC2 to CpG islands chromatin. The crystal structure of the N-terminal domains of PHF1 (PHF1- TPE) bound CpG-containing DNAs provides the first structural framework to demonstrate PCL proteins are crucial for PRC2 recruitment to CpG islands^26^. Our previous study showed that the PCL proteins stabilize the dimerization of PRC2 complexes and promotes PRC2-PCL targeting to CpG islands chromatin *in vivo*^21^. However, the mechanisms by which PRC2 is recruited to CpG islands chromatin are multifactorial and complex. Here, we expand we and other previous studies and further show PCLs can form biomolecular condensates through phase separation to facilitate PRC2 complex condensates formation on the CpG islands DNA. This work provides new perspective to show that PCL proteins link PRC2 to CpG islands via the previously uncharacterized phase-separation mechanism. Similar to the N-C intramolecular interaction of PCLs is sensitive to the methylated H3K36 (H3K36me3), the binding of CpG-DNA to EH domain also results in enhancing PCL droplets size and thus facilitating PCLs driven PRC2 condensates formation, suggesting the mechanism by PRC2 complex respond to the different chromatin environments to change the corresponding genes expression via its accessory factors PCLs phase separation, raising the question as to whether other PRC2 accessory factors may also evolve to use LLPS mechanism to regulate PRC2 condensate properties.

Phase separation play key roles in many cellular functions and dysregulation of biomolecular condensates formation is closely associated with many human disease ^47^. In this work, we observed aberrant PCL proteins droplet lost its chromatin colocalization and decreased PRC2 activity, indicating that the formation of an appropriate droplet size is necessary for PRC2 function includes enzymatic activity and chromatin recruitment. Actually, the similar phenomena were observed for various IDRs containing nuclear proteins, which phase separate into biomolecular condensates that mechanically exclude chromatin^37^. In the case of RNA regulates the size of phase-separated condensates further strengthens the concept that the optimum size of biomolecular condensates is critical for their biological functions^48, 49^. All of these indicate that condensation assembly and molecular escaping maintains an optimum physical condensate size as an important regulatory role for phase separation. Understanding of condensate function in normal and aberrant cellular states, and of the mechanisms of condensate formation, is providing new insights into human disease and revealing novel therapeutic opportunities^50^. Collectively, our studies show that the dysregulated expression of PCLs leads to the dysfunction of PRC2, including enzyme activity and chromatin dissociation, which may be closely associated with the abnormal phase-separated PCLs formation.

## Methods

### PHF1, MTF2, PHF19 and their mutant proteins expression and purification

Full- length PHF1, MTF2, PHF19 and their mutant proteins were both subcloned into a pET- 28a vector with an N-terminal His6-Sumo-mEGFP tag. The cDNA sequence encoding SUZ12(C2) domain (residues 146-363) was inserted into PGEX-4T1 vector and a TEV protease cleavage site was inserted between GST tag and SUZ12(C2). The wild-type or mutant His6-Sumo-EGFP-PCLs were co-expressed with the PGEX-4T1-TEV- SUZ12(C2) in *E.coli* BL21 (DE3) cells, respectively. The cells were grown in LB medium at 37°C to an OD600 of 0.8 and induced with 0.25mM IPTG at 20°C for 16 hr. The cells were lysed by sonication in lysis buffer containing 50mM Tris-HCl pH 8.0, 500mM NaCl, 5% glycerol and 5mM β-mercaptoethanol supplemented with 1mM PMSF. After centrifugation, the supernatant was applied to Ni-NTA resin. The protein bound to Ni-NTA resin was extensively washed using Ni-NTA wash buffer (50mM Tris pH 8.0, 500mM NaCl, 5% Glycerol and 5mM 2-mercaptoethanol) and then eluted with the wash buffer supplemented with 300 mM imidazole. The Strep-Tactin resin was used for further purification after the His6-Sumo tag was removed from EGFP-PCLs by SUMO protease. Protein eluted from the Strep-Tactin resin was pooled, flash frozen in liquid nitrogen and stored at -80°C.

### Reconstitution of mCherry-PRC2-PHF1 complex

Human mCherry-PRC2-PHF1 complex (EZH2-EED-SUZ12-RBBP4-PHF1) was co-expressed in Sf9 insect cells utilizing the Bac-to-Bac baculovirus expression system (Invitrogen). The mCherry- PRC2-PHF1 complex was purified essentially the same as previously described (Chen et al., 2018). Briefly, these constructs include Flag-EZH2, mCherry-Suz12, His6-Flag-Eed-3C-StrepII, mEGFP-PHF1-StrepII, and His6-Thrombin-Rbbp4. After 48 hr co-infection, Sf9 insect cells were harvested and washed twice with PBS prior to freezing at 80°C. For purification, cell pellets were resuspended in lysis buffer containing 50mM Tris, pH 8.0, 500mM NaCl, 1mM PMSF, 5mM β-mercaptoethanol and 5% glycerol and lysed by sonication. The complex was first purified by a Ni-NTA column, and then loaded onto a Strep-Tactin column (IBA) for further purification. The PRC2-PHF1 protein complex eluted from the Strep-Tactin column was concentrated, and stored at - 80°C. SDS-PAGE was used to analyze the complex eluted fractions.

### CpG-DNA synthesis and labeling

Different lengths of CpG islands DNA (CGI^Lhx6^) were amplified from mouse genomic DNA by PCR, which was used as previously described ^21^. The pacific blue (FAM-405) was labeled on the synthetic CGI^Lhx6^ DNA forward primers (CGI^Lhx6^ DNA-F). Through PCR amplification, we are able to obtain different lengths of FAM-405 labeled CGI^Lhx6^ DNA. The CGI^Lhx6^ DNA-F (5′- ATCCGCCCGCTCGACGCGCGCGCCGGGA-3′) and different reverse primers (CGI^Lhx6^ DNA (1-153)-R: TCCCGGCGCGCGTCGAGCGGGCGGAT; CGI^Lhx6^ DNA (1-353)-R: 5′-CATGTACTGGAAGCATGAGAGC-3′; CGI^Lhx6^ DNA (1-799)-R: 5′-AGTGGGTGAGCGTCGGGGATCCT-3′) were used in this study. The different lengths CGI^Lhx6^ DNA were obtained by PCR and further purified via a column and quantified with a ND-2000C NanoDrop spectrophotometer (NanoDrop Technologies). The 12 mer CpG-DNA with the palindromic sequence 5′-GGGCGGCCGCCC-3′ containing two CpG motifs was synthesized by Sangon Biotech (Shanghai), which were used in all pull-down experiments.

### CRISPR-Mediated PHF19 Knockout in Hela Cells

PHF19-KO Hela cell line was constructed by using the CRISPR/Cas9 gene editing system. Single-guide RNA (sgRNA) targeting sequence (5’-TGATCTTCCCGAGGTAGTAC-3’) was synthesized and cloned into the lentiCRISPR v2 expression vector. Lentiviruses was produced by using PSPAX2 and VSVG. Briefly, 2.25 μg PSPAX2, 0.75 μg VSVG, and 3 μg LentiCRISPR v2 sgRNAs for PHF19 construct were co-transfected into HEK-293T cells on a 60 mm cell culture dish by using PEI reagent. The Hela cells were then infected with the lentiviruses and selected with 2 μg/mL puromycin. Single-cell colonies were picked and the efficiency of the PHF19 knockout was assessed by western blot at protein levels and validated by DNA sequencing.

### Inducible ectopic protein expression in Hela cells

Lentiviruses were produced by co-transfecting 293T cells with PSPAX2, VSVG, and the PLVX-TetOne-BSD vectors as outlined in the Method Details. Subsequently, 3XFlag-PHF19-FL, 3XFlag-PHF19- TPE, 3XFlag-PHF19-IDR, 3XFlag-PHF19-RC, and 3XFlag-PHF19-ΔIDR knock-in Hela cell lines were generated by infecting PHF19-KO Hela cells with these lentiviruses, followed by selection using 3 μg/mL Blasticidin S (BSD) for one week. After the BSD selection, the cells were treated with a series of different concentration of Doxycycline (Dox). The inducible ectopic 3XFlag tagged PHF19 wild-type and mutants proteins expression were confirmed by western blot. Cells preparing for RNA sequencing, the aforementioned Hela cells, which include untreated Hela cells control, PHF19-KO Hela cells, rescued PHF19-KO Hela cells with a 3XFlag-PHF19(FL) cDNA, and rescued PHF19-KO Hela cells with a 3XFlag-PHF19-ΔIDR cDNA were cultured in a medium supplemented with 20 ng/mL Doxycycline (Dox) for 36 hours, and then harvest cells for RNA sequencing.

### *In vitro* phase separation experiment

All recombinant proteins were kept in stocking buffer (50 mM Tris pH 8.0, 500 mM NaCl and 5mM β-mercaptoethanol), which were thawed at room temperature and centrifuged at 15,000 rpm for 5 min to remove aggregates. Phase separation generation was induced at room temperature by diluting proteins and NaCl to the indicated concentrations. Otherwise indicated in the figure legend, the final concentration of NaCl was 150 mM for *in vitro* phase separation assays. In addition, the different concentrations of H3K36me3 or CpG-DNA were supplemented to induce phase separation of PCL proteins. After mixing, 10 µL of the reaction solution were loaded onto glass slides with coverslips and imaged using a Leica TCS SP8 microscope equipped with oil immersion objectives of 63× or 100×. Fiji software was used to prepare the images and quantify the size and number of droplets.

### Fluorescence recovery after photobleaching (FRAP) measurements

The FRAP experiments were performed on a Zeiss LSM 880 microscope equipped with a 100× oil immersion objective at room temperature. The mCherry tagged PCLs plasmid were transfected into HeLa cells, and biomolecular condensates were observed after 24 hr. A defined region of 1 µm diameter was photobleached by the equipped 561-nm laser pulse (3 repetitions, 90% intensity, dwell time of 5 second). The FRAP Profiler plugin for FIJI was used to measure the intensity of fluorescent signals at the bleached spot, the unbleached control spot as background. Fluorescence intensities of the region has been controlled by subtracting the background intensity from the unbleached spot. There are three unbleached (control) or three bleached (experimental) granules in each sample, with a mean and standard deviation of fluorescence intensities.

### Co-immunoprecipitation (Co-IP) and Western blot (WB)

For Co-IP, the expression plasmids were transfected into HEK293T cells (ATCC). After 24 hr, cultured cells were harvested and lysed with lysis buffer (25 mM Tris-HCl, pH 7.4, 150 mM NaCl, 1% Triton X-100, 1mM EDTA) supplemented with protease and phosphatase inhibitors for 45 min on ice. Cell lysate was then centrifuged at 10,000 rpm for 15 min at 4℃ to collect supernatant. The supernatant was incubated with GFP-Nanoab-Agarose Beads (LABLEAD , GNA-25-500) for 4 hr at 4℃. For GFP-tag pull-down, the supernatant was incubated with GFP-tagged recombinant proteins and GFP-Nanoab-Agarose Beads for 6 hr at 4℃. Next, beads were washed with lysis buffer and then boiled in SDS loading buffer (50 mM Tris-HCl pH 7.4, 2% SDS, 6% glycerol, 0.004% bromophenol blue, 1% β-mercaptoethanol) as samples for western blot analysis. Samples for western blot analysis were run on 5-10% SDS-polyacrylamide gel electrophoresis (SDS-PAGE) and transferred to nitrocellulose. After blocking in 5% milk at room temperature for one hour, the membranes were incubated with primary antibodies overnight at 4°C. Following washing with TBST, the membranes were incubated with secondary antibodies for 1 hour at room temperature, followed by washing. In order to visualize the immunoreactive products, enhanced chemiluminescence reagents and autoradiography were used.

### Dual-Luciferase Reporter Assay

The reporter assays were carried out as previously described ^28, 38^. For the PCLs and its mutants luciferase reporter assay, the PHF1 and it mutants (Including △TPE, △IDR and △RC)ΔMTF2 and it mutants (Including △TPE, △IDR and △RC), PHF19 and it mutants (Including △TPE, △IDR Δ △RC) DNA fragments were cloned into pBind-GAL4-DBD vector. The pBind-GAL4-DBD empty vector was employed as a control. We co-transfected 200 ng of the above interested GAL4-DBD plasmid with 200 ng of reporter vector with 6×GAL4UAS-TK-luciferase (G6-TK-luc) into 293T cells, respectively. We harvested cells from a cell culture, suspended them, and lysed them in 1× Reporter Lysis Buffer (Promega). Based on the manufacturer’s instructions, the activity of luciferase was measured using the Luciferase Assay System Kit (Promega). Using Renilla luciferase activity as a reference, transfection efficiency was calculated. Luminescence intensities of firefly luciferase were normalized to that of Renilla luciferase. Subsequently, the luminescence intensities were normalized to the control group. Three replicates were used.

### Determination of fraction condensed by sedimentation assay

The mEGFP-PHF1 was mixed with different concentrations of H3(29-43)K36me3 or 12-mer-CpG DNA, and the mCherry-PRC2-PHF1 complex was mixed with 799bp CpG-DNA (CGI^Lhx6^ DNA) and Histone octamer in 50 mM Tris and 150mM NaCl, pH 7.4. After incubation for 15 min at 25℃Δthe reaction mixture was centrifuged at 12,000 rpm for 15 min at 25℃. The absorbance of the supernatant was compared against the total concentration of respective fluorophores labelled proteins or DNA before the phase separation. The concentration of mEGFP-PHF1 or mCherry-PRC2-PHF1 was determined by measuring mEGFP absorbance at 488 nm with an extinction coefficient of 56,000 M^-1^cm^-1^ and mCherry absorbance at 587 nm with an extinction coefficient of 72,000 M^-1^cm^-1^, respectively.

### Immunofluorescence staining

In the experiments, mEGFP-, Flag-, or mCherry-labeled PCLs and its mutants were transfected into HeLa cells for 24 hr. After removing culture medium, cells were fixed in 4% paraformaldehyde (Sigma) for 15 min at room temperature and were washed with PBS for three times. Samples were permeabilized with PBS supplemented with 0.1% Triton-X 100 for 15 min at room temperature. Next, samples were incubated with primary antibody (see key resources table) overnight at 4°C. After washing for 10 min for three times, the samples were then incubated with secondary Alexa Flour antibody (1:200 dilution in PBS) for 2 hr at room temperature. In order to visualize the DNA, cells were stained with 40,6-diamidino-2-phenylindole (DAPI) for 10 min at room temperature and were washed with PBS for three times. Finally, we mounted slides with Slowfade Diamond Antifade reagent (Life Technologies) and acquired images using Zeiss LSM 880 (Zeiss, Germany) or Leica TCS SP8 (Leica Microsystems GmbH, Mannheim, Germany) confocal laser scanning microscopes. We were able to analyte the fluorescence signal by using the LAS-X viewer tool.

### Live cell imaging

HeLa Cells for live imaging were performed at a confluency of 50% on a 35 mm MatTek glass bottom dish (poly-D-lysine coated). Live cell imaging was carried out on a Zeiss 880 LSM with Airyscan, a planapochromatic 63× oil objective, and the Airyscan processing was carried out in ZEN software (Zeiss). We acquired a time-lapse movie of PCL proteins at 100 milliseconds per frame for 30 seconds, which was displayed as a kymograph. Images were analyzed using Zeiss ZEN software or ImageJ.

### Intrinsic disorder tendency analysis

In order to analyze the intrinsic disorder tendency of PHF1, MTF2, and PHF19, the Predictor of Natural Disordered Regions (PONDR) version SVL2 and VL3 (http://www.pondr.com/) was used. The scores were assigned between 0 and 1, and a score above 0.5 indicates that there is a disorder present.

### Statistical analysis and reproducibility

Pearson colocalization coefficients of different proteins were determined by ImageJ. The size and number of PCLs or PRC2 droplets were measured using ImageJ. The measurements were performed over the entire cell. An analysis of the data was conducted using the GraphPad Prism 8.0 program in order to conduct statistical analysis. Statistical significance was calculated from three or more groups by using one-way ANOVA with Tukey’s corrections for comparison of three or more groups, the two groups using unpaired two-tailed Student’s t-tests. In all cases, the data are presented as the mean values, the standard error of mean of at least three technical replicates. Unless otherwise stated, significance was defined as any statistical outcome that resulted in a P value of at least 0.05 in order to determine its significance.

### Reporting Summary

Further information on research design is available in the Nature Research Reporting Summary linked to this article.

### Data availability

All other relevant data supporting the key findings of this study are available within the article and its Supplementary Information files or from the corresponding author upon reasonable request. Source data are provided with this paper.

## Acknowledgements

This work was supported by the National Natural Science Foundation of China (32270638 to S.C., 32100464 to S.C., 82302954 to S.P., and 82000171 to F.H.), the Science, Technology and Innovation Commission of Shenzhen Municipality (JCYJ20230807091204009 to S.C.), the Natural Science Foundation of Fujian (2022J01051 to S.C. and 2020J05291 to F.H.), the Postdoctoral Science Foundation of China (2023M732956 to S.P.), and the Open Research funds for Fujian Provincial Key Laboratory of Innovative Drug Targets Research (FJ-YW-2022KF06 to S.P.), the Fundamental Research Funds for the Central Universities (20720220122 to S.C.) and the Nanqiang Outstanding Young Talents Program from Xiamen University. The graphical abstract was generated with BioRender.

## Author contributions

S.C. conceived the project. S.C. and S.P. designed and performed most of the experiments with help from H.X., B.Z., F.H., X.W., C.W., M.Z., X.L. and M.J. Q.C., X.G. and X.Z. contributed technical assistance. F.S. performed bioinformatic analyses. S.C. and S.P. wrote the manuscript.

