## supplemental Files for "Phase separation of Polycomb-like (PCL) proteins drive PRC2 complex condensates to regulate gene expression"

**This PDF file includes:**

**Figs. S1 to S7**

### Supplementary Figure 1. (Peng et al.)

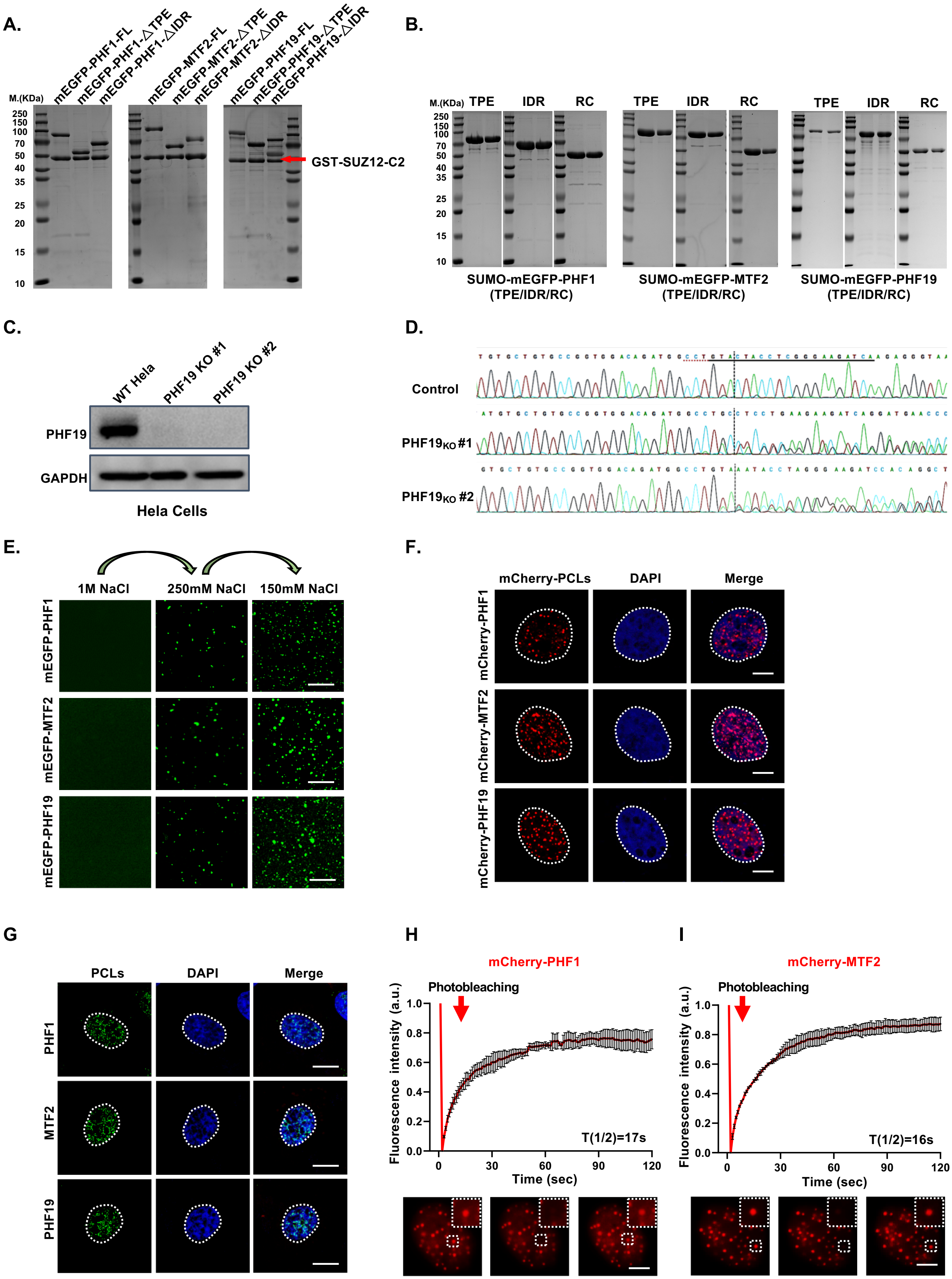

**Supplementary Figure 1. PCL proteins undergo phase separation in vitro and in vivo.**

**(A)** SDS-PAGE gel of the purified mGFP-PCLs-GST-SUZ12(C2) binary protein complexes.

**(B)** Analysis of PCLs and its mutants purified from *E.coil* was performed on SDS-PAGE and visualized using Coomassie blue staining.

**(C)** Western blotting showed that PHF19 was knocked out in HeLa cells.

**(D)** The PHF19 gene was knocked out by sequencing analysis.

**(E)** Representative images showing PCLs proteins condensates formed at indicative NaCl concentrations. Scale bar,10 μm.

**(F)** Fixed-cell images of mCherry-tagged PCL proteins in HeLa cells. Scale bars, 5 μm.

**(G)** Endogenous PCLs display droplets in HeLa cells revealed by immunostaining with corresponding antibody. Scale bar, 5 μm.

**(H-I)** FRAP curves and representative images of experiments with mCherry-PHF1 or mCherry-MTF2 droplets in HeLa cells. Data were presented as mean values ± SEM (n= 3 independent experiments). Scale bar, 5 μm.

### Supplementary Figure 2. (Peng et al.)

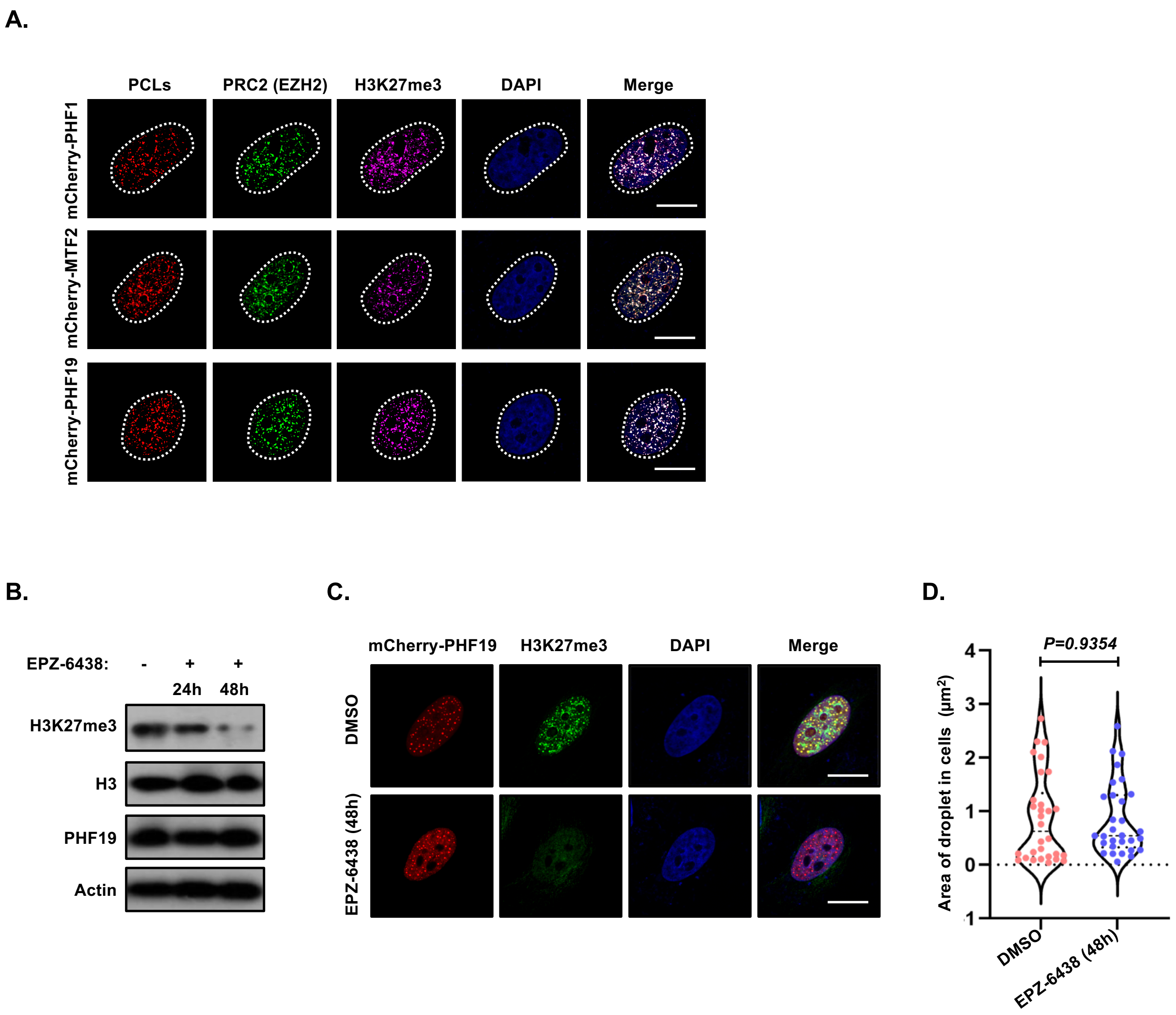

**Supplementary Figure 2. PCL proteins undergo phase separates with PRC2 in vivo.**

**(A)** Representative images showing colocalization of transfected mCherry-tagged PCLs with Flag-EZH2 and H3K27me3 in HeLa cells. Scale bars, 5 μm

**(B)** Western blot analysis showed that HeLa cells treated with EPZ-6438 (10 μM) could lead to a global loss of H3K27me3 levels.

**(C)** Representative images show the effect of EPZ-6438 on H3K27me3 expression and colocalization with mCherry-PHF19 droplets. Scale bars, 5μm.

**(D)** Quantitative analysis of mCherry-PHF19 forming droplets in HeLa cells after treatment with EPZ-6438 (10 μM, 48h). A two-tailed unpaired Student's t-test was used for statistical analysis, and data are presented as mean values ± SEM (n= 30 independent samples).

### Supplementary Figure 3. (Peng et al.)

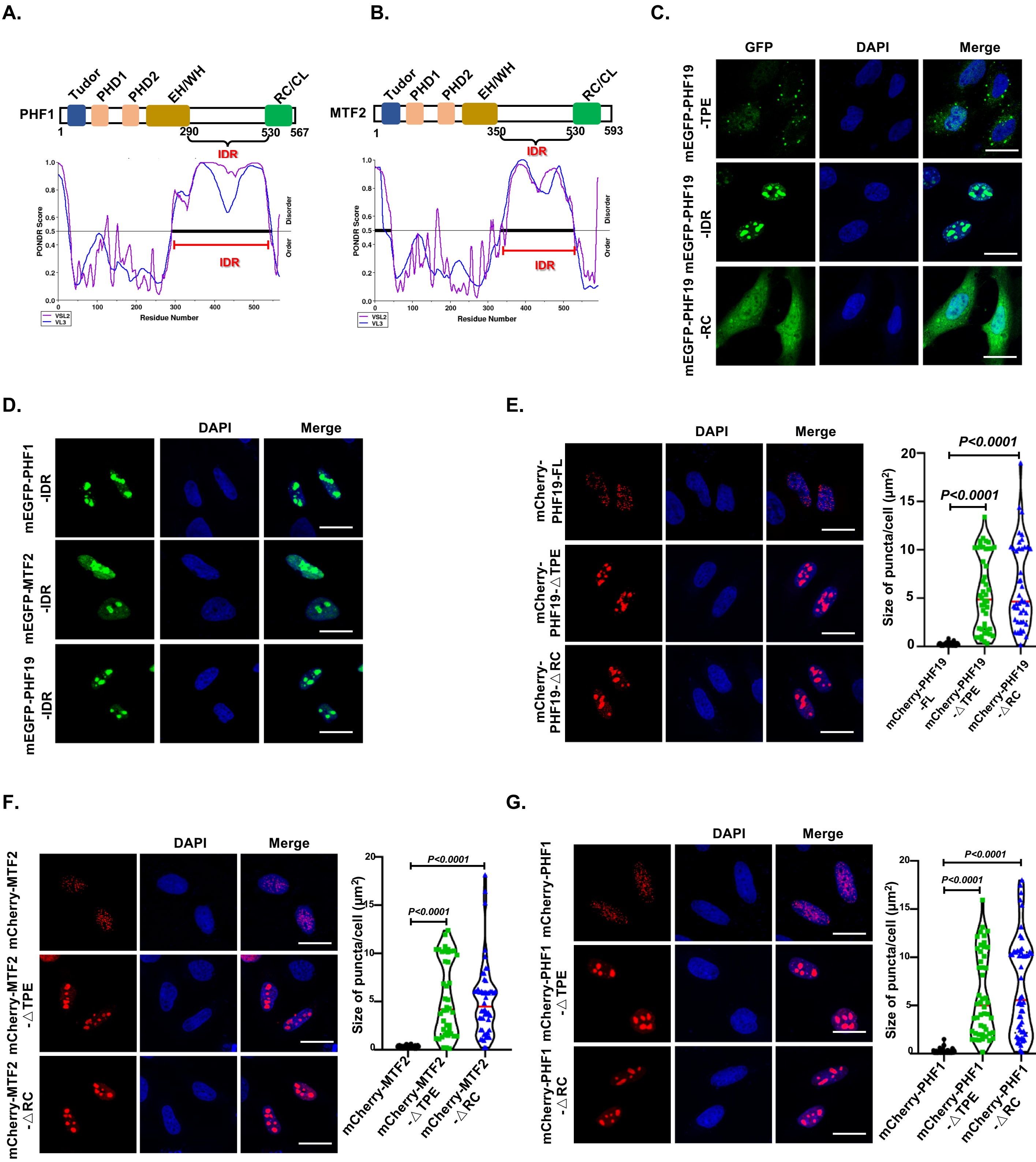

**Supplementary Figure 3. The IDR domain of PCL proteins is required for condensate formation.**

**(A)** Graph plotting intrinsic disorder with predictor of intrinsically disordered regions (PONDR) using the VSL2 and VL3 algorithm for PHF1. IDR, intrinsically disordered region.

**(B)** Graph plotting intrinsic disorder with predictor of intrinsically disordered regions (PONDR) using the VSL2 and VL3 algorithm for MTF2. IDR, intrinsically disordered region.

**(C)** Representative images showing mEGFP-PHF19 and its mutants droplets formation in HeLa cells. Scale bar, 10 μm.

**(D)** Representative images showing mEGFP-PCLs-IDR droplets formation in HeLa cells. Scale bar, 10 μm.

**(E-G)** Representative images and quantitative analysis showing the size of PCL proteins formation droplets compared with indicated mutants. The one-way ANOVA was used for statistical analysis, and data are presented as mean values ± SEM (n = 50 independent samples). Scale bar, 10 μm.

### Supplementary Figure 4. (Peng et al.)

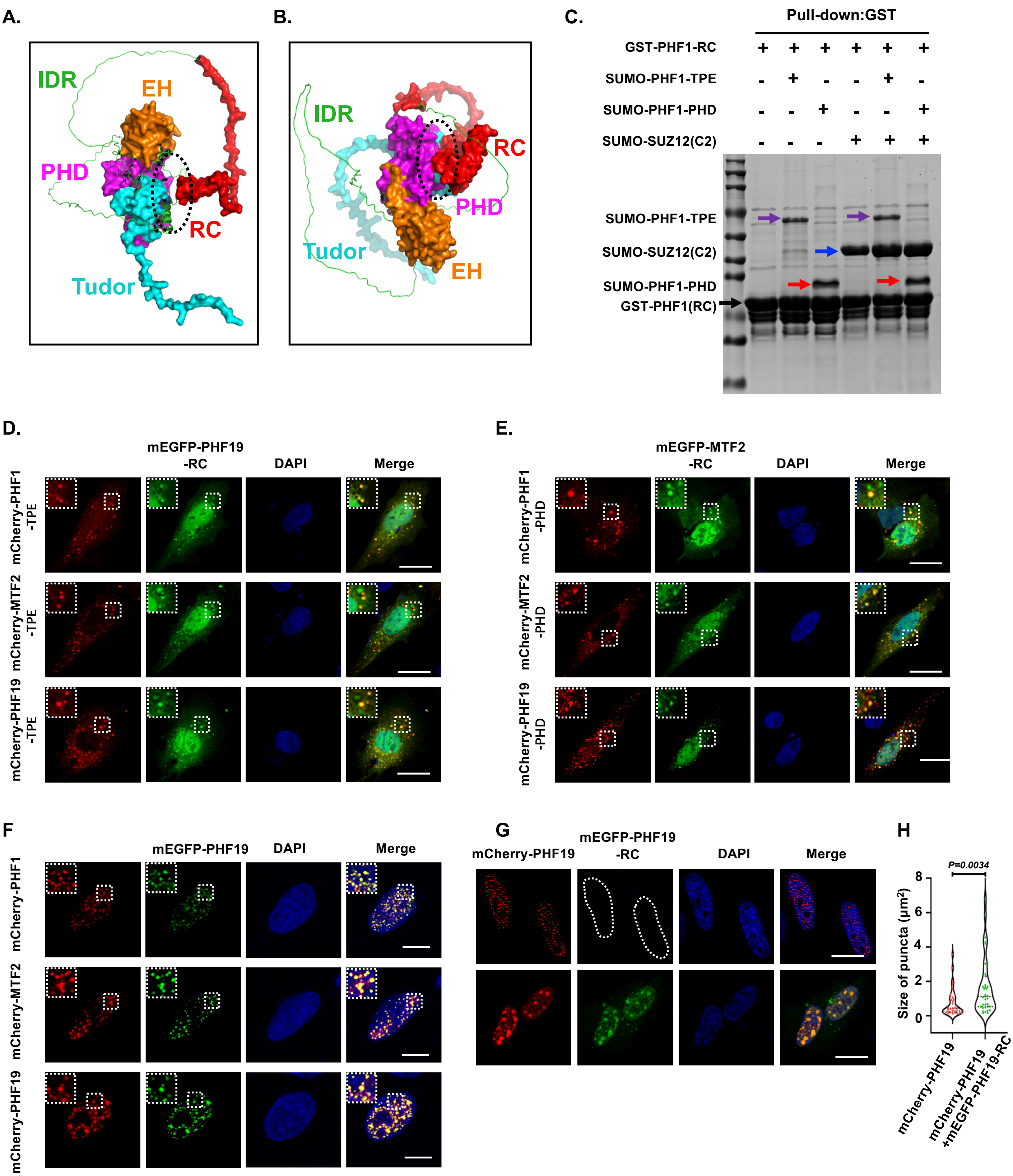

**Supplementary Figure 4. Intramolecular interaction regulates the phase separation of PCL proteins.**

**(A-B)** The structure of PHF1 and MTF2 were predicted using the AlphaFold algorithm. The black dotted line indicates the protein interaction interface.

**(C)** GST-Pull down examine recombinant GST-PHF1-RC protein was interaction with SUMO-PHF1-TPE or SUMO-PHF1-PHD in the absence or presence with SUMO-SUZ12(C2).

**(D)** Representative images show colocalization of transfected mEGFP-PHF19-RC with mCherry-PHF1-TPE, mCherry-MTF2-TPE or mCherry-PHF19-TPE in HeLa cells. Scale bars, 10 μm.

**(E)** Representative images show colocalization of transfected mEGFP-MTF2-RC with mCherry-PHF1-PHD, mCherry-MTF2-PHD, or mCherry-PHF19-PHD in HeLa cells. Scale bars, 10 μm.

**(F)** Representative images show colocalization of transfected mEGFP-PHF19 with mCherry-PHF1, mCherry-MTF2, or mCherry-PHF19 in HeLa cells. Scale bars, 5 μm.

**(G-H)** Representative images and quantitative analysis show the effect of mEGFP-PHF19-RC overexpression on droplet size of mCherry-PHF19 in cells. A two-tailed unpaired Student's t-test was used for statistical analysis, and data are presented as mean values ± SEM (n = 50 independent samples). Scale bars, 10 μm.

### Supplementary Figure 5. (Peng et al.)

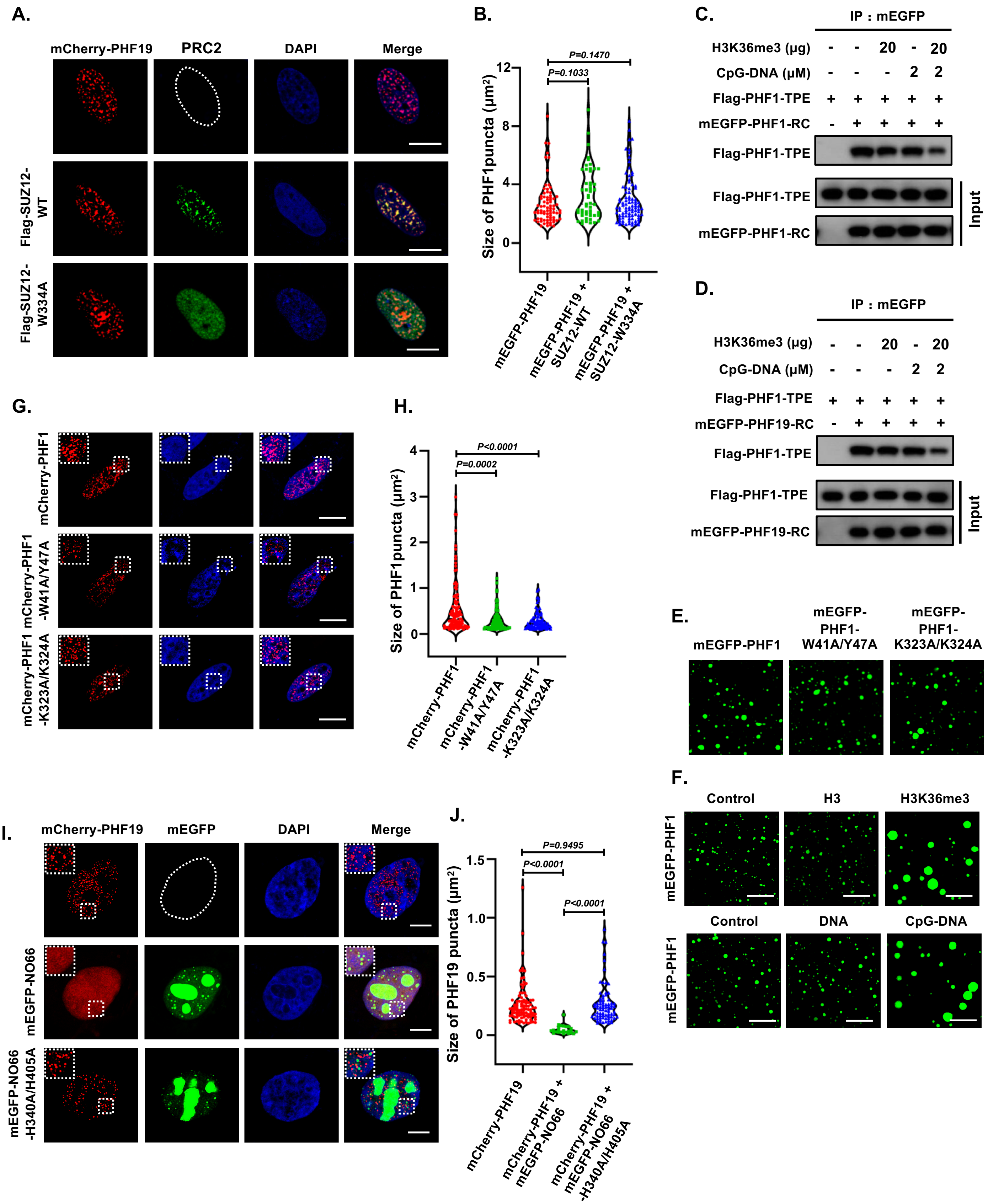

**Supplementary Figure 5. Effect of H3K36me3 and CpG islands on phase separation of PCL proteins.**

**(A-B)** Representative images and quantitative analysis show the effect of Flag-SUZ12 and Flag-SUZ12$^{W334A}$ over-expression on mCherry-PHF19 droplets formation in cells. The one-way ANOVA was used for statistical analysis, and data are presented as mean values ± SEM (n= 50 independent samples). Scale bar, 5 μm.

**(C)** Effect of H3K36me3 and CpG-DNA on the interaction between Flag-PHF1-TPE and mEGFP-PHF1-RC *in vitro*.

**(D)** Effect of H3K36me3 and CpG-DNA on the interaction between Flag-PHF1-TPE and mEGFP-PHF19-RC *in vitro*.

**(E )** The representative picture show the phase separation of PHF1 and its mutants *in vitro*. Scale bar, 10 μm.

(F) Representative images show that CpG-DNA or H3K36me3 promotes mEGFP-PHF1 droplets formation, but not ordinary DNA or H3 peptide. Scale bar, 10 μm.

**(G-H)** Representative images and quantitative analysis show the effect of W41A/Y47A and K323A/K324A point mutation on PHF1 droplets formation *in vivo*. The one-way ANOVA was used for statistical analysis, and data are presented as mean values ± SEM (n= 50 independent samples). Scale bar, 5 μm.

**(I-J)** Representative images and quantitative analysis showed that mEGFP-NO66 over-expression inhibited mCherry-PHF19 phase separation in cells, but not mEGFP-NO66-$^{H340A/H405A}$ mutants. The one-way ANOVA was used for statistical analysis, and data are presented as mean values ± SEM (n= 50 independent samples). Scale bar, 5 μm.

### Supplementary Figure 6. (Peng et al.)

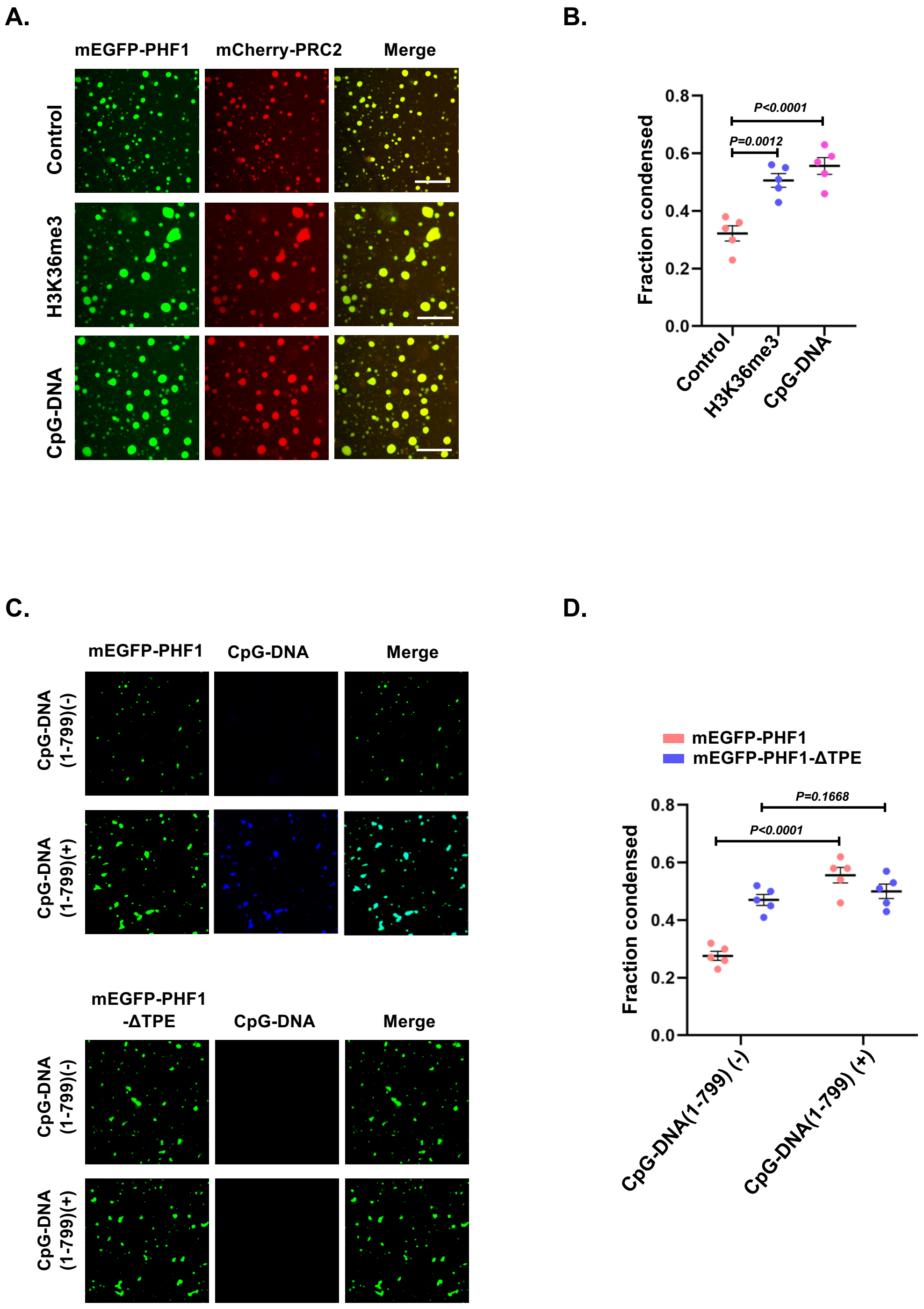

**Supplementary Figure 6. PHF1 phase separation promotes the condensation of PRC2 to CpG-DNA chromatin.**

**(A-B)** Representative images and quantitative analysis show the effect of CpG-DNA and H3K36me3 peptide on PRC2-PHF1 complex phase separation. The fraction of condensed PRC2-PHF1 complex as a function of CpG-DNA or H3K36me3 peptide. The one-way ANOVA was used for statistical analysis, and data are presented as mean values ± SEM (n= 5 independent samples). Scale bars, 10 μm.

**(C-D)** Representative images and quantitative analysis show that CpG-DNA promotes mEGFP-PHF1 phase separation depends on n-terminal TPE. The fraction of condensed mEGFP-PHF1 and its mutants as a function of CpG-DNA. The one-way ANOVA was used for statistical analysis, and data are presented as mean values ± SEM (n= 5 independent samples). Scale bars, 10 μm.

### Supplementary Figure 7. (Peng et al.)

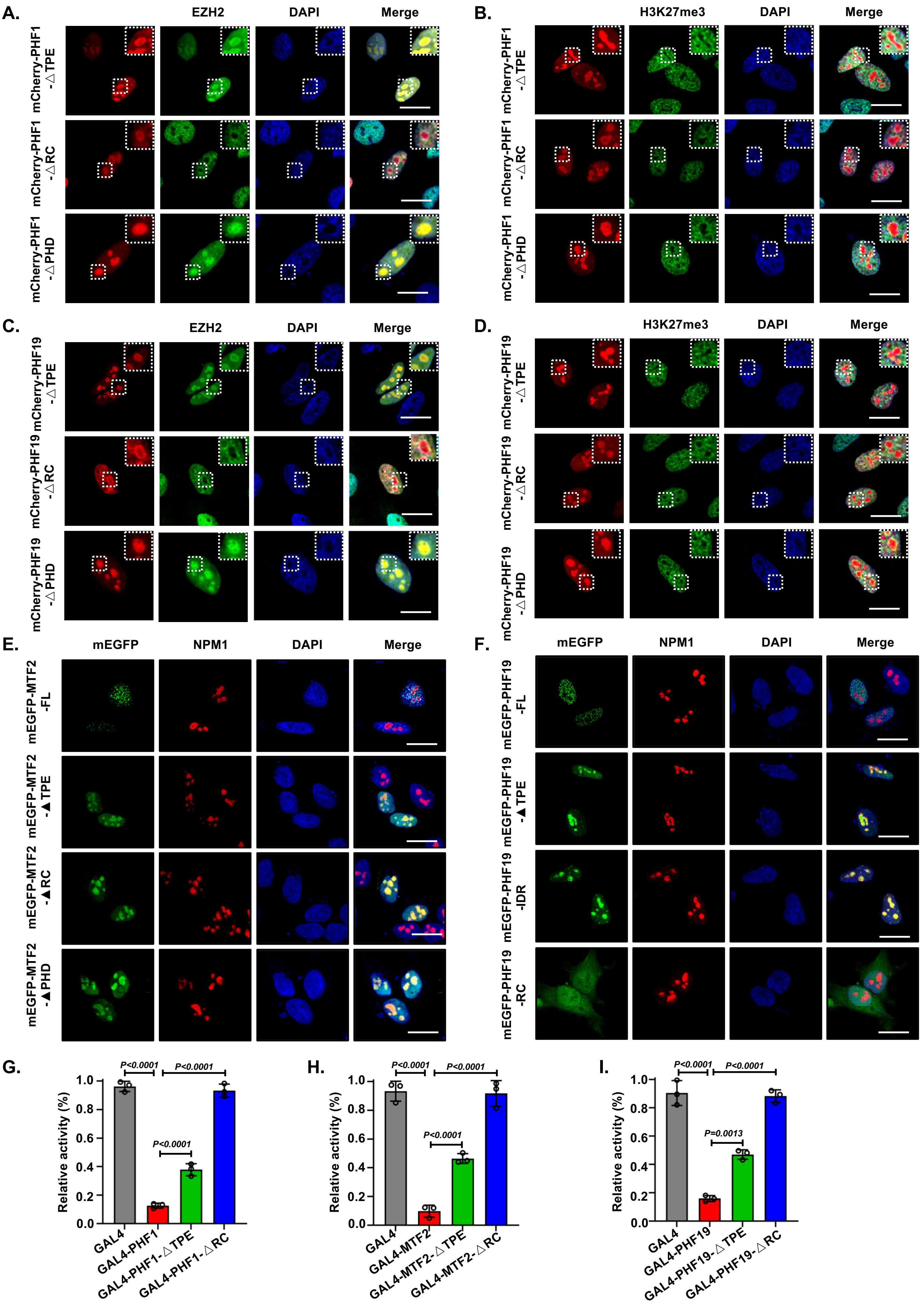

**Supplementary Figure 7. PCLs phase separation regulates PRC2 localization chromatin and transcriptional repression.**

**(A-B)** Representative images show colocalization of transfected mCherry-PHF1 and its mutants with endogenous EZH2, H3K27me3 and chromatin marked by DAPI in HeLa cells. Scale bars, 10 μm.

**(C-D)** Representative images show colocalization of transfected mCherry-PHF19 and its mutants with endogenous EZH2, H3K27me3 and chromatin marked by DAPI in HeLa cells. Scale bars, 10 μm.

**(E)** Representative images show MTF2 and its mutants co-localizing with nucleolar marker NPM1. Scale bars, 10 μm.

**(F)** Representative images show PHF19 and its mutants co-localizing with nucleolar marker NPM1. Scale bars, 10 μm.

**(G-I)** Luciferase gene repression of PCLs and its mutants. The one-way ANOVA was used for statistical analysis, and data are presented as mean values ± SEM (n= 3 independent experiments).
